# Neural and muscular contributions to the age-related loss in power of the knee extensors in men and women

**DOI:** 10.1101/2023.10.24.563851

**Authors:** David J. Wrucke, Andrew Kuplic, Mitchell Adam, Sandra K. Hunter, Christopher W. Sundberg

**Author notes:** **Correspondence:** Corresponding Author Christopher W. Sundberg, Ph.D. Department of Physical Therapy Marquette University Cramer Hall Room 215 604 North 16^th^ Street Milwaukee, WI 53233.

## Abstract

The mechanisms for the loss in limb muscle power in old (60-79 years) and very old (≥80 years) adults and whether the mechanisms differ between men and women are not well-understood. We compared maximal power of the knee extensor muscles between young, old, and very old men and women and identified the neural and muscular factors contributing to the age-related loss of power. 31 young (22.9±3.0 years, 15 women), 83 old (70.4±4.9 years, 39 women), and 16 very old adults (85.8±4.2 years, 9 women) performed maximal isokinetic contractions at 14 different velocities (30- 450°/s) to identify peak power. Voluntary activation (VA) and contractile properties were assessed with transcranial magnetic stimulation to the motor cortex and electrical stimulation of the femoral nerve. The age-related loss in power was ∼6.5 W·year^-1^ for men (*R*^2^=0.62, *p*<0.001), which was a greater rate of decline (*p*=0.002) than the ∼4.2 W·year^-1^ for women (*R*^2^=0.77, *p*<0.001). Contractile properties were the most closely associated variables with power output for both sexes, such as the rate of torque development of the potentiated twitch (men: *R*^2^=0.69, *p*<0.001; women: *R*^2^=0.57, *p*<0.001). VA was weakly associated with power in women (*R*^2^=0.13, *p*=0.012) but not men (*p*=0.191), whereas neuromuscular activation (EMG amplitude) during the maximal power contraction was not associated with power in men (*p*=0.347) or women (*p*=0.106). These data suggest that the age-related loss in power of the knee extensor muscles is due primarily to factors within the muscle for both sexes, although neural factors may play a minor role in older women.

**NEW & NOTEWORTHY:** The accelerated age-related loss in power relative to the loss in muscle mass of the knee extensors was primarily due to factors altering the contractile properties of the muscle for both old and very old (≥80 yr) adults. The mechanisms for the decrements in power with aging appear largely similar for men and women, although neural factors may play more of a role in older women.

## INTRODUCTION

Human aging is accompanied by a progressive decline in the neuromuscular system that ultimately leads to decreased physical function and quality of life for older adults. Mechanical power output of limb muscle, the product of force or torque and velocity, has emerged as one of the strongest predictors for the decrements in physical function with aging (1), with the decline in power typically beginning around the fourth to fifth decade of life and accelerating thereafter (2-4). Importantly, the age-related decrements in power output vastly exceed the age-related loss in total muscle mass (2, 3, 5, 6), suggesting other factors, such as a reduced ability of the nervous system to activate the muscle and/or factors disrupting the contractile properties of the muscle, must also be contributing to the age-related loss in power. Despite the importance of limb muscle power output to physical function in older adults, the extent that neural and muscular mechanisms contribute to the accelerated age-related decline in power relative to muscle mass remains poorly understood.

Several impairments in the neuromuscular system may contribute to the accelerated loss in power output relative to muscle mass with aging, including, a reduced ability of the nervous system to voluntarily activate the muscle (7-11), instability of the neuromuscular junction (12), impaired cross-bridge mechanics and Ca^2+^ handling (13-15), and/or the selective atrophy of fibers expressing the fast myosin heavy chain (MyHC) II isoforms (16-18). Continued advancements in noninvasive stimulation procedures can be used to help localize the primary sites along the motor pathway where age-related impairments occur and may contribute to power loss. For example, neural drive from the motor cortex during maximal voluntary contractions can be estimated with the interpolated twitch technique by delivering single magnetic pulses to the cortical motor neurons via transcranial magnetic stimulations (TMS) (19-21). In addition, the contractile properties of the muscle can be evaluated with TMS by measuring the involuntary relaxation rates following the depolarization elicited by the magnetic pulse (22, 23) or by measuring the kinetics of the potentiated resting twitch elicited by supramaximal electrical stimulations to the peripheral nervous system. Limitations with current stimulation paradigms, however, restrict measurements of voluntary activation with an interpolated stimulus to isometric or slow velocity contractions (24); thus, integrating measures of surface electromyography (EMG) obtained during the moderate-to high-velocity contractions required to elicit peak power along with measures of TMS and electrical stimulation will help localize the primary sites along the motor pathway contributing to power loss with aging.

Identifying the primary mechanisms for the accelerated age-related loss in power relative to muscle mass is particularly important in cohorts that may be most affected by the detrimental effects of losing muscle power, such as older women and very old adults (≥80 yr). For example, young adult women generate less power than men before the decrements that occur from aging (2, 3, 25, 26), which may result in older women being more susceptible to having insufficient power to maintain mobility and perform activities of daily living (2, 27-29). Both cross-sectional and longitudinal studies have observed that the rates of absolute power loss with aging are greater in men than women, but do not differ between the sexes when expressed relative to muscle mass or as a percent decline (2-4). These findings suggest that the mechanisms for the age-related loss in power may be similar between men and women. However, studies investigating potential sex differences in the decrements in power with aging have used a limited number of loads and velocities (2-4, 30, 31), rather than obtaining peak power across the full range of the torque-velocity relationship. This is important because aging is known to slow the shortening velocity of whole muscle (31-33), which would likely result in peak power occurring at a wide range of velocities across young, old and very old men and women. Furthermore, no studies have performed a comprehensive assessment of the torque-velocity relationship and coupled the assessment with measures of surface EMG, TMS, and electrical stimulation to determine if the primary sites along the motor pathway contributing to the age-related loss in power differs between men and women.

In addition to potential sex-based differences, the declines in neuromuscular function and skeletal muscle atrophy accelerate with advancing age in men and women ≥80 yrs old (2, 5, 34, 35). The mechanisms for the accelerated decline in neuromuscular function of very old adults, particularly power output, are unknown, but it has been postulated from a study on the knee extensor muscles of very old women (85-92 yrs) that the ability of the nervous system to voluntarily activate the muscle optimally may become impaired, with the problem exacerbated in immobile women (36, 37). No age differences, however, were observed in the ability to voluntarily active the knee extensors in very old men when assessed with TMS to the motor cortex (25) or electrical stimulation to the femoral nerve (38), suggesting the impairments in activation may be specific to very old women. The extent that impairments in neural activation are contributing to power loss in very old adults is unknown. Moreover, it was recently observed that the decrements in power output of the lower extremity in men and women aged 80 years of age were both 48% of the sex-matched young cohorts (2), which suggests that the mechanisms of power loss might be similar for very old men and women.

Thus, the purpose of this study was to identify the primary neural and muscular factors contributing to the loss in peak power output of the knee extensor muscles with advanced age in old (60-79 yr) and very old (≥80 yr) men and women. We hypothesized that the absolute rates of power loss across the age groups would be greater in men than women, but the rate would be similar between the sexes when power is normalized to muscle mass or expressed relative to the sex-matched young adults. We also hypothesized that factors altering the contractile properties within the muscle would be the most closely associated with the decrements in peak power for the old and very old men and women and that neural factors would only be modestly associated with power in the very old adults and/or older women.

## METHODS

### Participants and Ethical Approval

One hundred and thirty individuals participated in this study: 31 young (19-33 years, 16 men and 15 women), 83 old (61-79 years, 44 men and 39 women), and 16 very old adults (80-93 years, 7 men and 9 women). Participants provided written informed consent and underwent a general health screening that included a questionnaire where older participants were required to score ≥26 on the mini mental state (39) to participate in the study. Participants were healthy, community-dwelling adults free of any known neurological, musculoskeletal, and cardiovascular diseases. All experimental procedures were approved by the Marquette University Institutional Review Board and conformed to the principles in the Declaration of Helsinki. Anthropometrics and physical activity levels for the participants are reported in Table 1.

**Table 1:** Anthropometrics and physical activity levels for the young, old, and very old men and women.

| Variable | Units | Young ($\leq 35$ yrs) | | Old (60-79 yrs) | | Very Old ($\geq 80$ yrs) | |
| --- | --- | --- | --- | --- | --- | --- | --- |
|  |  | Men (16) | Women (15) | Men (44) | Women (39) | Men (7) | Women (9) |
| Age*‡ | yrs | 23.3 ± 3.6 | 22.6 ± 2.2 | 70.3 ± 4.2 | 70.5 ± 5.5 | 89.2 ± 3.4 | 83.1 ± 2.6 |
| Height*† | cm | 176.6 ± 8.3 | 164.6 ± 5.5 | 177.1 ± 7.9 | 162.1 ± 4.9 | 167.6 ± 5.4 | 160.5 ± 6.3 |
| Weight† | kg | 76.1 ± 10.5 | 65.1 ± 9.6 | 83.0 ± 13.3 | 68.6 ± 15.0 | 71.9 ± 4.4 | 71.9 ± 11.8 |
| BMI* | kg·m<sup>-2</sup> | 24.0 ± 1.7 | 24.0 ± 3.2 | 26.7 ± 3.8 | 26.1 ± 5.5 | 25.6 ± 1.6 | 27.9 ± 4.4 |
| Body Fat*† | % | 18.2 ± 3.6 | 29.6 ± 7.3 | 29.7 ± 6.6 | 39.1 ± 7.4 | 30.0 ± 6.8 | 39.8 ± 6.7 |
| Whole Body Lean Mass†‡ | kg | 60.3 ± 9.5 | 43.4 ± 4.2 | 55.5 ± 7.1 | 39.3 ± 4.8 | 48.4 ± 3.8 | 42.9 ± 7.9 |
| Thigh Lean Tissue Mass*†‡ | kg | 7.2 ± 1.4 | 5.3 ± 0.7 | 6.5 ± 1.0 | 4.5 ± 0.7 | 5.2 ± 0.4 | 4.8 ± 0.6 |
| Physical Activity* | steps·day<sup>-1</sup> | 8623 ± 3973 (12) | 9825 ± 2840 (14) | 8494 ± 4657 (36) | 7437 ± 3664 (30) | 4134 ± 3283 | 3455 ± 1643 |

Body fat percentage was measured via dual X-ray absorptiometry (Lunar iDXA, GE, Madison, WI). Physical activity was measured via triaxial accelerometry (GT3X, ActiGraph, Pensacola, FL). The sample sizes (*n*) for each cohort and certain variables are reported in parentheses. Symbols next to the variable name indicate a significant effect of \* age, † sex, or an ‡ age x sex interaction at $p \leq 0.05$ . Values are reported as means ± SD.

### Experimental Protocol and Setup

The experimental protocol to measure the neuromuscular performance of the knee extensor muscles consisted of 1) 3-5 maximal voluntary isometric contractions (MVC) performed without stimulations, 2) an assessment of torque and power outputs during maximal concentric isokinetic contractions across a range of 14 different velocities, and 3) 5 sets of isometric contractions at torques equivalent to 100%, 60%, and 80% of the MVC coupled with transcranial magnetic stimulation (TMS) and femoral nerve stimulation to assess voluntary activation and involuntary contractile properties of the muscle (Fig. 1). The experimental setup to measure the neuromuscular performance of the knee extensors was identical to the setup described previously (25). Briefly, testing was performed on the dominant leg of each participant (preferred kicking leg) except when the participant reported a previous surgical procedure, knee or leg pain, or osteoarthritis of the dominant leg (1 young woman, 2 old women, 3 very old women, 2 old men, and 1 very old man). Participants were seated upright in the high Fowler’s position with the starting knee position set at 90° flexion in a Biodex System 4 Dynamometer (Biodex Medical, Shirley, NY) (Fig. 1). The position of the dynamometer was adjusted so that the axis of rotation of the dynamometer’s lever arm was aligned with the axis of rotation of the participant’s knee. The length of the dynamometer’s lever arm was adjusted for each participant and secured with a Velcro strap proximal to the malleoli. Extraneous movements and changes in the hip angle were minimized by securing the participants to the seat with the dynamometer’s four-point restraint system. To ensure the measured torques and velocities were generated primarily by the knee extensor muscles, participants were prohibited from grasping the dynamometer with their hands.

**Fig. 1.**
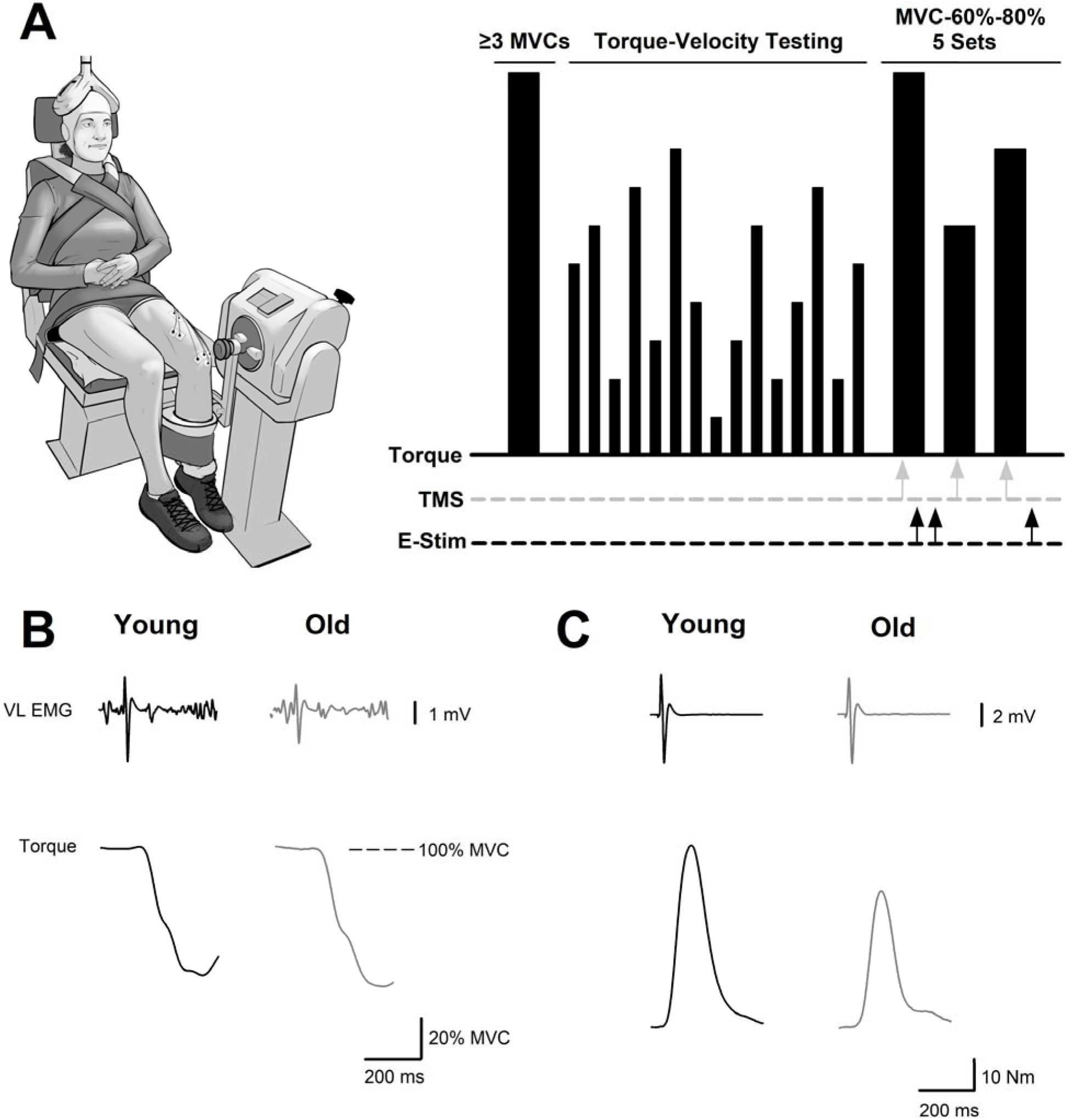
Experimental setup and protocol. (A) Experimental setup of the research participants (borrowed with permission from Sundberg *et al*., 2018) and schematic of the experimental protocol to measure the neural and muscular factors associated with the age-related power loss in men and women. Participants performed a minimum of 3 knee extensor maximal voluntary isometric contractions (MVCs) with no stimulations followed by assessments of maximal torque and power output across a range of 14 different isokinetic velocities. The session concluded with five sets of isometric contractions that were comprised of a MVC followed by contractions at both 60 and 80% MVC (MVC-60-80%). TMS to the motor cortex and electrical stimulation to the femoral nerve during the MVC-60-80% contractions are represented by the gray and black arrows, respectively. The x-axis for the experimental protocol in panel (A) is not to scale, and the timing of the stimuli and contractions are described in detail in the METHODS. (B) Representative data from a young and older man showing a motor evoked potential (MEP) from the vastus lateralis and a superimposed twitch (SIT) elicited by TMS to the motor cortex along with (C) a compound muscle action potential from the vastus lateralis and a potentiated resting twitch (Q_tw_) elicited by electrical stimulation to the femoral nerve.

### Isokinetic Contractions

Each participant performed 15 sets of 4 maximal isokinetic knee extension contractions across a range of 14 different velocities (Fig. 2). The range of motion (ROM) was set at 95° to minimize the additional braking force applied by the dynamometer near the end of the ROM, and the test velocities were 30, 60, 90, 120, 150, 180, 210, 240, 270, 300, 330, 360, 400, and 450°/s. Strong verbal encouragement was provided to kick as hard and as fast as possible throughout the full range of motion for each velocity. Participants were required to take a minimum of two minutes rest after each set of 4 contractions to minimize fatiguing the muscle during the torque-velocity assessment. To objectively evaluate if fatigue occurred, the first and last trials were performed at the same 180°/s velocity. Power outputs in the last set of 180°/s contractions were 3 ± 11% higher than in the first 180°/s contractions (*p* = 0.013) indicating no evidence of muscle fatigue during the torque-velocity assessment but a mild “warm-up effect’ may have occurred. Univariate analysis of variance showed there was no effect of age (*p* = 0.062, 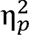 = 0.03) or sex (*p* = 0.503, 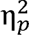 = 0.00) on the difference in power between the first and last sets of 180°/s contractions. To minimize the influence of an order effect, the remaining 13 velocities were randomized to make 5 different sequences of isokinetic velocities. Each participant was then randomly assigned to 1 of the 5 sequences for their experimental assessment.

**Fig. 2.**
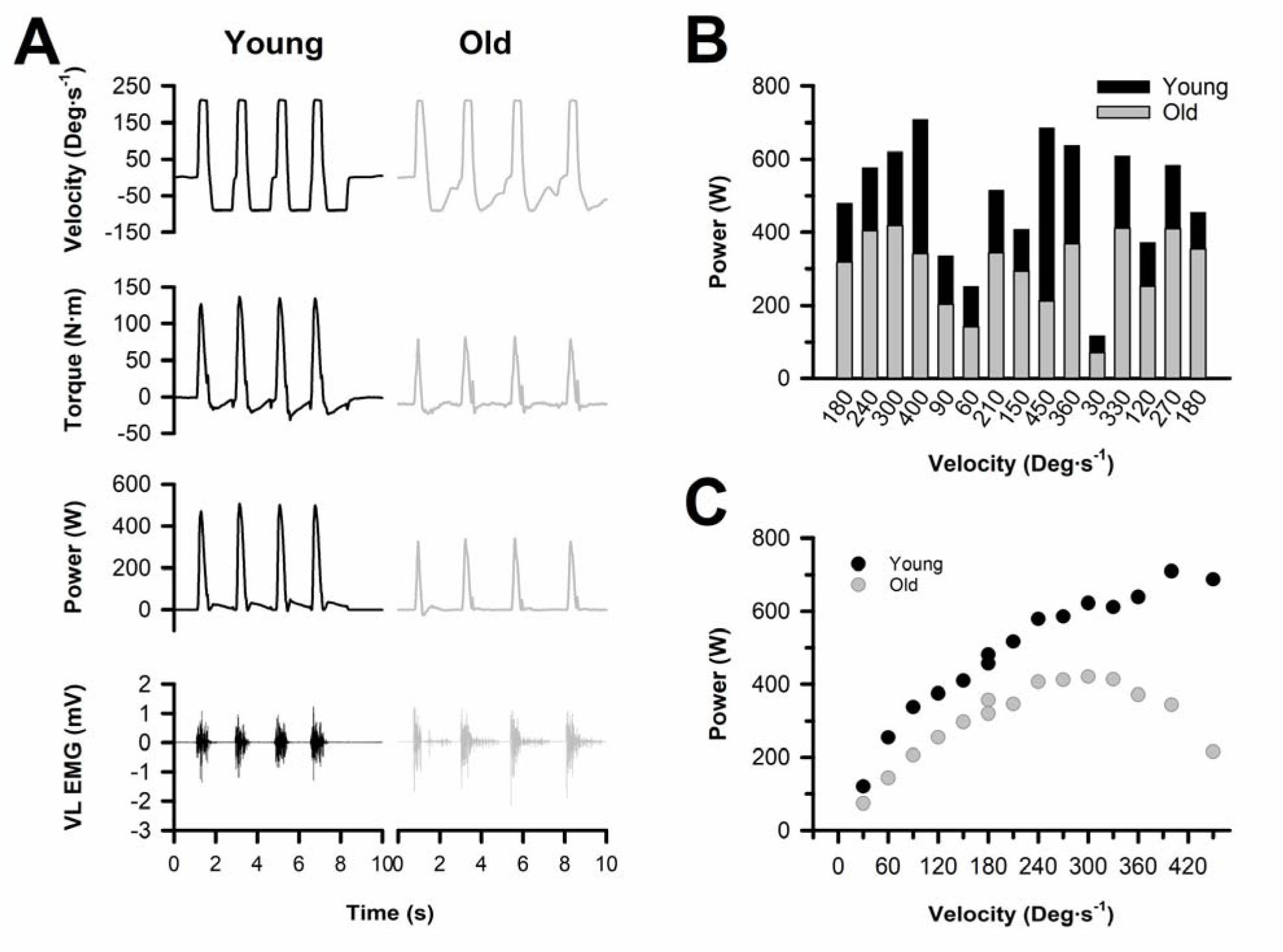
Representative isokinetic torque-velocity data. (A) Representative velocity, torque, power, and vastus lateralis EMG recordings from the 7^th^ set of 4 contractions performed at 210°/s by a young and older man. (B) Peak power output at the 14 different velocities (30 – 450°/s) shown in the random order for one of the testing sequences. Participants were randomly assigned to complete one of five randomized isokinetic velocity sequences. All subjects started and ended with contractions performed at 180°/s to test whether fatigue occurred during the protocol. (C) Power-velocity curves for the 14 different isokinetic velocities (30 – 450°/s) from the young and older man.

### MVC-Stim Protocol

The experiment began with electrical stimulation of the femoral nerve to identify the electrode placement that elicited the maximum peak-to-peak compound muscle action potential (maximum M-wave: M_max_) of the vastus lateralis (VL), rectus femoris (RF) and vastus medialis (VM). Following the electrical stimulations, participants performed a minimum of 3 brief (2-3 s) knee extension MVCs without stimulation. Participants were provided strong verbal encouragement and visual feedback on their performance with a 56 cm monitor mounted 1-1.5 m directly in front of their line of vision. Each MVC was interspersed with at least 60 s rest, and MVC attempts were continued until the two highest values were within 5% of each other. The highest torque output from the MVCs was used to calculate 1) the target forces for the submaximal isometric contractions needed for optimizing the TMS parameters (i.e., coil placement and stimulator intensity) and 2) the visual feedback gain in the subsequent MVC trials used to assess voluntary activation.

Once the optimal TMS position and intensity was identified following the torque-velocity assessment with the isokinetic contractions, participants performed five sets of brief isometric contractions (2-3 s per contraction) with the knee extensor muscles to obtain the measures used in identifying the mechanisms of power production. Each set of contractions included a MVC followed by contractions at 60% and 80% MVC (MVC-60-80%) with TMS delivered at each contraction to estimate the resting twitch amplitude for the calculation of voluntary activation (19, 40). Single-pulse femoral nerve stimulation was delivered during the MVC and at rest immediately following (<5 s) both the MVC and 80% MVC contractions. Sets of MVC-60-80% contractions were interspersed with at least 2.5 min rest to help ensure repeatable maximal efforts were performed while minimizing residual fatigue from each set. The median value from the sets of MVC-60-80% contractions was reported and used for the association analyses with peak power production of the knee extensors.

### Measurements and Data Analysis

#### Torque and Mechanical Power Output

Torque, position, and angular velocity from the dynamometer’s transducers were digitized at 500 Hz with a Power 1401 A/D converter and stored online using Spike 2 software [Cambridge Electronics Design (CED), Cambridge, UK]. The torque during each MVC was quantified as the average value over a 0.5 s interval centered on the peak torque of the contraction. The MVC value for each participant was the greatest isometric torque output recorded during the experimental session. Power outputs during the isokinetic assessments were quantified as the product of the instantaneous torques and angular velocities during the concentric contraction. Peak power for each isokinetic velocity was reported as the highest power output during only the portion of the contraction where the target velocity was achieved (Fig. 2).

#### Surface Electromyography (EMG)

Surface Ag/AgCl EMG electrodes (Grass Products, Natus Neurology, Warwich, RI) were adhered to the skin in a bipolar arrangement overlying the muscle bellies of the vastus lateralis, vastus medialis, rectus femoris, and biceps femoris with an inter-electrode distance of 2.5 cm. The skin was shaved and cleaned with 70% ethanol prior to electrode placement, and the reference electrodes were placed on the patella. Analog EMG signals were amplified (100 X), filtered (13-1,000 Hz band pass, Coulbourn Instruments, Allentown, PA), and digitized at 2,000 Hz with a Power 1401 A/D converter and stored online using Spike 2 software (CED). To best capture the EMG activity that elicited peak power, EMG during the isokinetic assessments were analyzed in two ways: 1) as the average RMS EMG from the onset of power production until 45 degrees of knee extension and 2) from the onset of power production until peak power at each target velocity was achieved. The results from the two methods did not differ, so only the average RMS EMG from the onset of power production until peak power generation are reported.

#### Electrical Stimulation

The femoral nerve was stimulated with a constant-current, variable high-voltage stimulator (DS7AH, Digitimer, Welwyn Garden City, Hertforshire, UK) to obtain M_max_ of the vastus lateralis, vastus medialis and rectus femoris. The cathode was placed over the nerve high in the femoral triangle, and the anode was placed over the greater trochanter. Single 200-µs square-wave pulses were delivered with a stimulus intensity beginning at 50 mA and increased incrementally by 50-100 mA until both the unpotentiated resting twitch torque amplitude and M_max_ for all three quadriceps muscles no longer increased. The intensity was then increased by an additional 20% to ensure the stimuli were supramaximal (range 120-720 mA).

Contractile properties of the knee extensor muscles were quantified with the potentiated resting twitches from the single-pulse femoral nerve stimulations delivered after the MVC and 80% MVC contractions (Fig. 1). Stimuli were delivered after both the MVC and 80% MVC contractions to ensure that at least one of the stimuli was delivered while the participant was fully relaxed. The values for each participant were the median obtained from the five sets of MVC-60-80% performed and were reported for the amplitude of the potentiated resting twitch torque (Q_tw_: N·m), the half relaxation time (ms), and the peak rate of torque development (Nm·s^-1^). The peak rate of torque development was quantified with the derivative of the torque channel as the highest rate of torque increase over a 10 ms interval.

#### Transcranial Magnetic Stimulation (TMS) & Voluntary Activation

Optimal stimulator position and intensity were identified with the method described previously (25). Briefly, the motor cortex was stimulated by delivering a 1-ms duration magnetic pulse with a concave double-cone coil (110 mm diameter: maximum output 1.4 T) connected to a monophasic magnetic stimulator (Magstim 200^2^, Magstim, Whitland, UK). The optimal stimulator position was determined as the location that elicited the greatest motor evoked potential (MEP) in the vastus lateralis while the subject contracted at 20% MVC. This position was marked to ensure repeatable placement of the coil for the remainder of the experiment. The stimulator intensity for the voluntary activation measurements was identified during brief (2-4 s) isometric contractions at 40% MVC. Single pulse TMS was delivered during each contraction with an intensity starting at 50% stimulator output and increased incrementally by 10% until the peak-to-peak MEP amplitude of the vastus lateralis failed to increase further or began to decrease. If the latter occurred, then the stimulator output was reduced in 5% decrements until the largest peak-to-peak MEP amplitude was achieved in the vastus lateralis. The intensity eliciting the largest MEP was compared to the intensity eliciting the largest twitch torque at the 40% MVC to verify that the stimulator intensities were approximately similar. This additional step ensured that the stimulus intensity did not elicit large activation of the antagonist muscles. This method was used instead of quantifying the biceps femoris MEP amplitude (%M_max_) because of the difficulty of maximally stimulating the sciatic nerve with surface electrical stimulation. It should also be noted that we’ve previously reported that the knee flexor MVC was on average only ∼40% of the knee extensor MVC at the 90° knee flexed position (25). Thus, the effect of any inadvertent activation of the antagonist muscle group on measurements of voluntary activation would be diminished due to the positioning of the participant.

Voluntary activation was quantified from each set of MVC-60-80% contractions based on the technique originally developed for the elbow flexors (19) and later for the knee extensors (21). Briefly, single pulse TMS was delivered during the MVC, 60% and 80% MVC contractions, and the amplitude of the superimposed twitch torque measured for each contraction. A linear regression was performed between the superimposed twitch torque and the voluntary torque to obtain an estimated resting twitch (eRT) by extrapolating the regression to the y-intercept. The resting twitch evoked by TMS was estimated rather than measured directly, because the excitability of the corticospinal tract increases markedly from rest to maximal activation (41). Any 3-point regression with an R^2^ < 0.8 (40) was excluded from the voluntary activation calculations. This occurred in 8.6% of the MVC-60- 80% sets. We were unable to obtain estimated resting twitches from one old woman, and an additional six old women and two young men did not complete the MVC-60-80% protocol due to the discomfort of the stimulations. It should also be noted that recent criticisms of the validity of the estimated resting twitch in the knee extensors have prompted a new method of calculating voluntary activation using the superimposed twitch from TMS and the resting twitch from femoral nerve stimulation (42). As a result, to provide comparison of the data to all other studies that have used TMS for voluntary activation and to align with emerging methods, we quantified voluntary activation for each set of MVC-60-80% contractions in two ways:

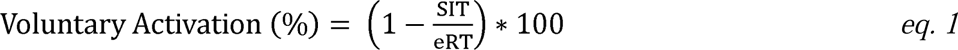

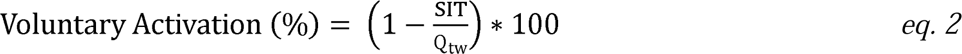

where SIT is the amplitude of the superimposed twitch torque elicited by TMS during the MVC, eRT is the calculated estimated resting twitch torque, and Q_tw_ is the torque amplitude of the potentiated resting twitch elicited by femoral nerve stimulation. The reported voluntary activation for each participant was the median from the 5 MVC-60-80% sets.

Absolute (Nm·s^-1^) and normalized (s^-1^) peak rates of torque relaxation were also determined from the TMS delivered during the MVC contractions (23). When TMS is delivered to the motor cortex during an MVC, there is a brief transient withdrawal of the descending neural drive following the stimulus that causes the muscle to involuntarily relax. The peak rate of torque relaxation was quantified with the derivative of the torque channel as the greatest rate of torque decrease over a 10 ms interval.

#### Thigh Lean Tissue Mass

Body composition and thigh lean tissue mass were assessed by dual X-ray absorptiometry (Lunar iDXA; GE, Madison, WI, USA). Thigh lean tissue mass was quantified for the region of interest using the manufacturer’s software (enCORE, version 14.10.022; GE), with the distal demarcation set at the tibiofemoral joint and the proximal demarcation set as a diagonal bifurcation through the femoral neck. DXA measures of thigh lean mass with these landmarks are strongly correlated with measures from magnetic resonance imaging but underestimate the age-related loss in thigh muscle mass (43).

#### Physical Activity Assessment

Physical activity was quantified for each participant with a triaxial accelerometer (GT3X, ActiGraph, Pensacola, FL) worn around the waist for at least 4 days (2 weekdays and 2 weekend days) as done previously (44). Data were reported for each participant as long as the accelerometer was worn for a minimum of 3 days (45).

#### Functional Performance Tests

The 14 tasks of the Berg Balance Scale were performed and scored as described previously (46). For the 6-minute walk test, subjects were asked to walk as far as possible along a 30 m minimally trafficked corridor for a period of 6 min with the primary outcome measure being the 6-min walk distance measured in meters (47). For the stair climb test, subjects were asked to climb 10 stairs as quickly as possible, and the time to complete the task was recorded (48). For the sit-to-stand test, the time for subjects to sit and stand 5 times and 10 times from a standard chair was recorded (49).

### Statistical Analyses

Simple linear regression analyses were performed between measurements of strength and power and age to estimate the rate of strength and power loss with aging. Individual two-factor univariate analyses of variance (ANOVA) were performed for the participant characteristics and mechanistic variables with age (young, old, or very old) and sex (men or women) as the grouping variables, and Bonferroni post-hoc tests were used to make pairwise comparisons between the age groups when a main effect of age was observed. Because the sample sizes decreased at the higher velocities of the isokinetic trials (Fig. 3), we analyzed the torque-velocity data with a linear mixed-model approach, with the isokinetic velocity used as the repeated measure (15 levels: 30 – 450°/s). The dependent variables for the mixed-model design were torque, normalized torque, power, and EMG, and the fixed effects were isokinetic velocity (15 levels: 30 – 450°/s), age group (3 levels: young, old, or very old), and sex (2 levels: men or women). Simple linear regression analyses were performed between peak power production and the neuromuscular stimulation measurements to identify the neural and muscular contributions to the age-related decrease in limb power production.

**Fig. 3.**
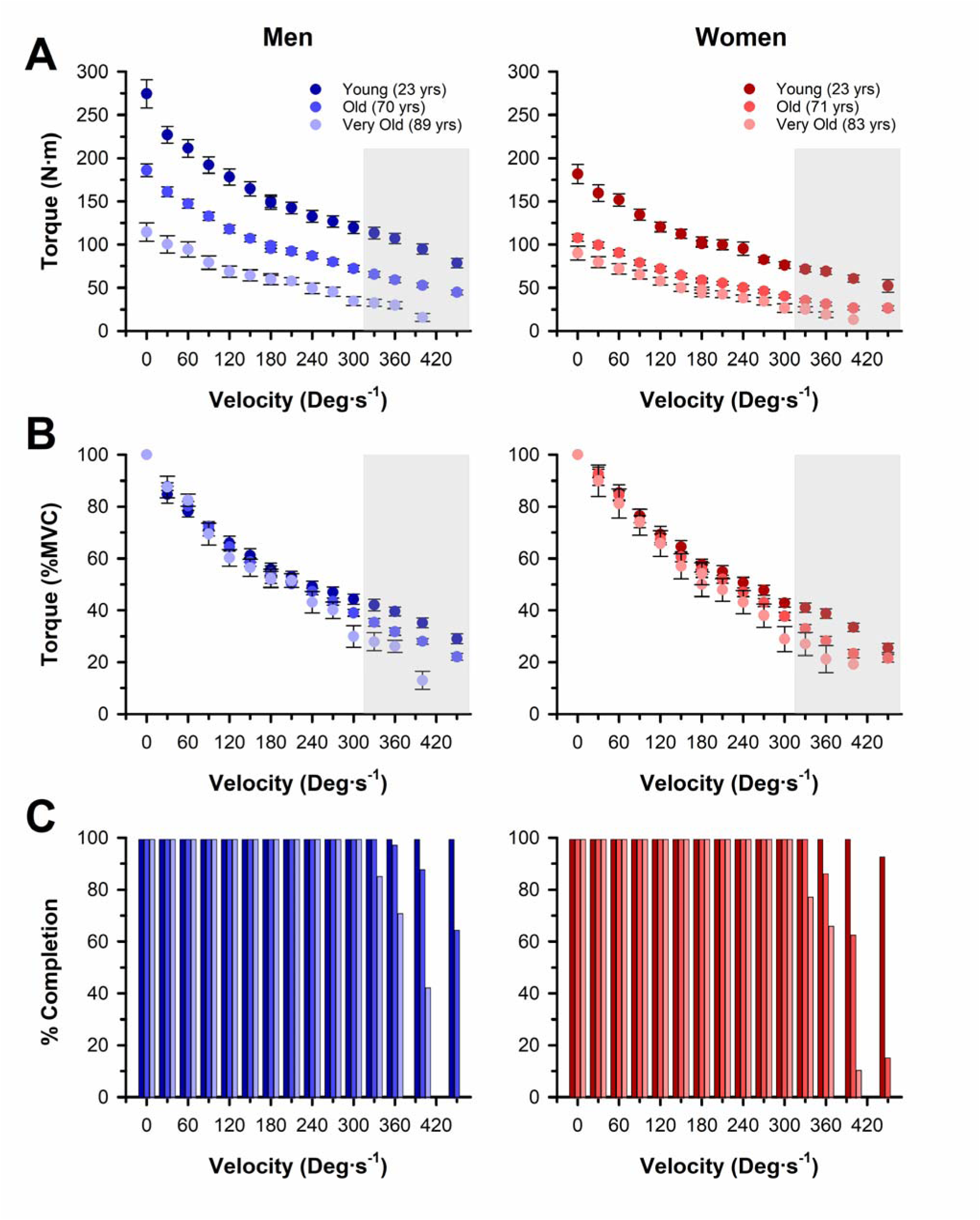
Maximal torque output across a range of isokinetic velocities. (A) Young adults produced more absolute torque than old and very old adults (*p* < 0.001) with a greater age difference at faster contraction velocities (velocity x age effect, *p* < 0.001). Men produced more absolute torque than women (*p* < 0.001) with a greater sex difference at faster contraction velocities (velocity x sex effect, *p* < 0.001). (B) Younger adults produced more torque relative to the MVC torque than old and very old adults (*p* < 0.001) with a greater age difference at faster contraction velocities (velocity x age effect, *p* < 0.001); however, the relative torque did not differ between men and women (*p* = 0.177). (C) Percentage of participants that achieved each tested isokinetic velocity. All subjects achieved velocities up to and including 300°/s. Gray boxes in panels (A) and (B) indicate velocities with <100% completion. Values are the means ± SEM.

Normal distributions and homogeneity of variance of the data were tested prior to any statistical comparisons with the Kolmogorov–Smirnov test and Levene’s statistic, respectively. If the assumptions of a normal distribution and/or homogeneity of variance were violated, then the non-parametric Kruskal-Wallis test was performed instead of the univariate ANOVA, with age and/or sex as the grouping variables, and Mann-Whitney U tests were used to make pairwise comparisons between the age groups when a main effect of age was observed. Due to the inability of the Kruskal-Wallis test to identify age x sex interactions, two-factor univariate analyses of variance (ANOVA) were performed to identify age x sex interactions regardless of violated assumptions of normality and homogeneity of variance. Interactions were only reported if confirmed by an age x sex interaction from univariate analyses of covariance (ANCOVA) using age as a continuous variable.

All significance levels were set at p ≤ 0.05 and all statistics were performed using SPSS (version 28, IBM, Chicago, IL). Data are presented as the mean ± standard deviation (SD) in the text and tables and the mean ± standard error of the mean (SE) in the figures. Effect sizes are reported as 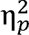 for main effects and Cohen’s *d* for post hoc pairwise comparisons.

## RESULTS

### MVC Torque Output

MVC torque outputs are reported in Table 2. MVC torque output for men decreased with age (*p* < 0.001, 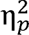 = 0.44) and was 38% and 61% lower in the old (*p* < 0.001, *d* = 1.51) and very old (*p* < 0.001, *d* = 2.49) compared with the young, and 37% lower in the very old compared with the old (*p* = 0.002, *d* = 0.91). For women, MVC torque also decreased with age (*p* < 0.001, 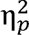 = 0.47) and was 44% and 57% lower in the old (*p* < 0.001, *d* = 1.80) and very old (*p* < 0.001, *d* = 2.88) compared with the young and 24% lower in the very old than the old (*p* = 0.027, *d* = 0.67). These decrements across age groups corresponded with an age-related loss in MVC torque of ∼2.2 N·m·year^-1^ for men (*R*^2^ = 0.55, *p* < 0.001), which was a greater rate of decline (*p* = 0.026, 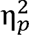 = 0.04) than the ∼1.5 N·m·year^-1^ for women (*R*^2^ = 0.57, *p* < 0.001). In contrast, when the MVC torques were expressed relative to the mean of the young men and women, the relative rate of decline did not differ between the men (∼0.9 %·year^-1^) and women (∼0.9 %·year^-1^, *p* = 0.978, 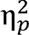 = 0.00). When we restricted the regression analyses to include only the older adults (old and very old groups, n = 99, age range 61-93 yrs.), the age-related loss in MVC torque was ∼3.4 N·m·year^-1^ for men (*R*^2^ = 0.30, *p* < 0.001), which was a greater rate (*p* = 0.027, 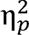 = 0.05) than the ∼1.3 N·m·year^-1^ in women (*R*^2^ = 0.11, *p* = 0.020). The relative rate of decline in MVC torque of the old and very old adults did not differ in the men (∼1.4 %·year^-1^) compared with women (∼0.8 %·year^-1^, *p* = 0.163, 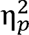 = 0.02).

**Table 2:**
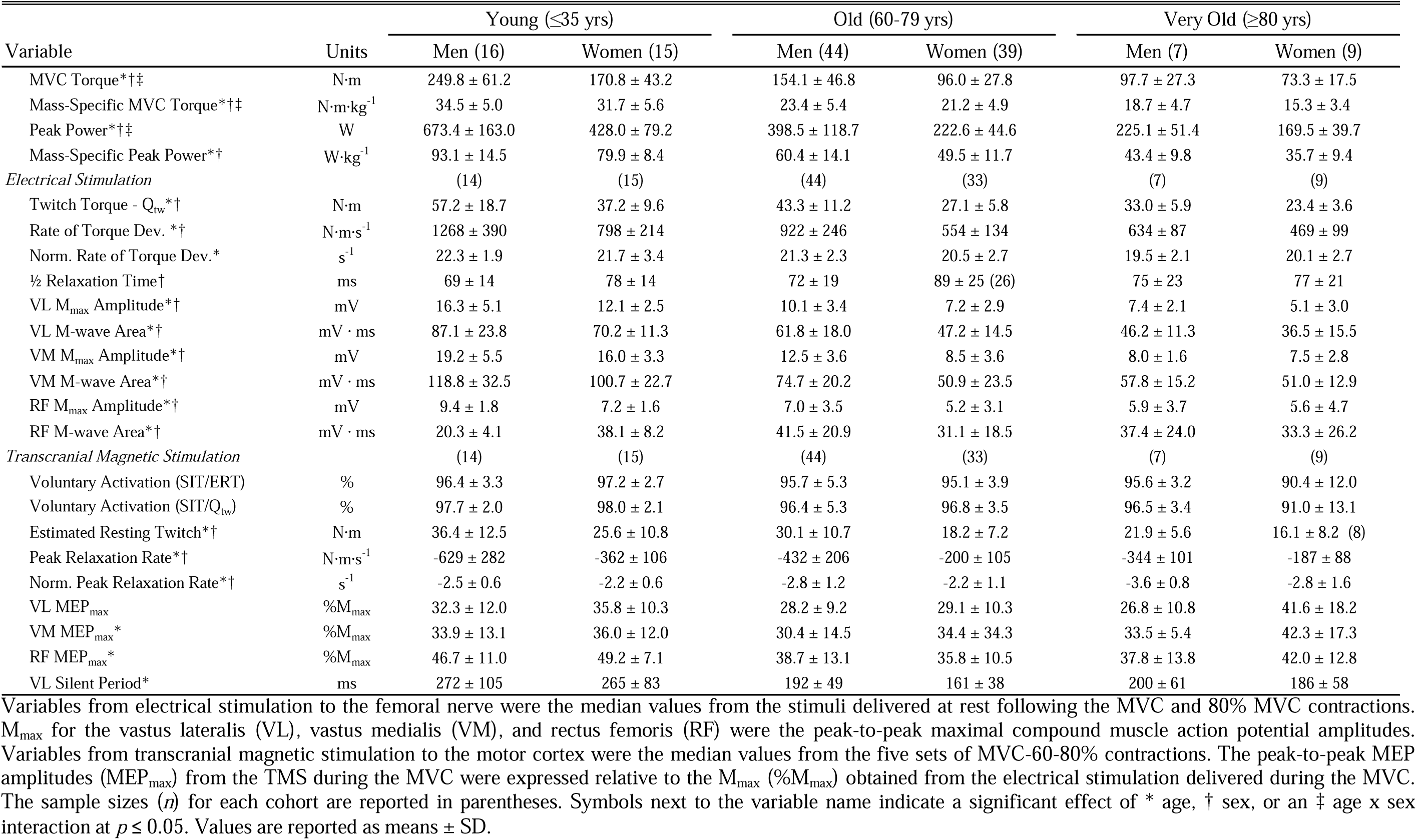
Neuromuscular performance measures from the young, old, and very old men and women.

### Torque-Velocity Relationship

All participants were able to successfully attain contraction velocities ≤300 °/s, but several participants were unable to achieve the contraction velocities >300 °/s (Fig. 3). The number of contractions successfully performed at velocities >300°/s differed with age (*p* < 0.001, 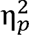 = 0.41) and was greater in the young than the old (*p* < 0.001, *d* = 0.50) and very old (*p* < 0.001, *d* = 1.34) and greater in the old than the very old (*p* < 0.001, *d* = 0.65). Men also successfully attained more high velocity contractions than women irrespective of age (*p* = 0.011, 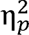 = 0.05).

Absolute torque outputs across the entire torque-velocity relationship differed with age (*p* < 0.001, 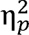 = 0.59) and were greater in the young than the old (*p* < 0.001, *d* = 3.57) and very old adults (*p* < 0.001, *d* = 5.40), and greater in the old than the very old adults (*p* < 0.001, *d* = 1.40, Fig. 3). The age differences in torque were greater at slower velocities (velocity x age effect, *p* < 0.001, 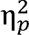 = 0.26). Men produced higher torques compared to women irrespective of age (*p* < 0.001, 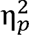 = 0.29) with a greater difference at slower velocities (velocity x sex effect, *p* < 0.001, 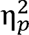 = 0.20) and a greater sex difference among the younger compared to the older groups (age x sex effect, *p* < 0.001, 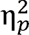 = 0.06). As expected, all groups had decrements in torque with increasing velocities (velocity effect, *p* < 0.001, 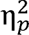 = 0.90).

To account for the differences in absolute maximal torque between the groups, the torques at each isokinetic velocity were expressed relative to the subject’s MVC. The relative torques differed with age (*p* < 0.001, 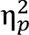 = 0.05) and were greater in the young than the old (*p* < 0.001, *d* = 0.60) and very old adults (*p* < 0.001, *d* = 0.88) with a greater age difference at higher velocities (velocity x age effect, *p* = 0.004, 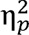 = 0.18, Fig. 3). There were no sex differences in relative torque production (*p* = 0.177, 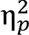 = 0.00), and all groups, irrespective of age or sex, had decrements in relative torque with increasing velocities (velocity effect, *p* < 0.001, 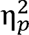 = 0.95).

### Power-Velocity Relationship

#### Power Output

Absolute power outputs across the power-velocity relationship differed with age (*p* < 0.001, 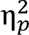 = 0.70) and were greater in the young than the old (*p* < 0.001, *d* = 3.16) and very old (*p* < 0.001, *d* = 4.70) and greater in the old than the very old (*p* < 0.001, *d* = 1.24). The age differences in absolute power were greater at faster velocities (velocity x age effect, *p* < 0.001, 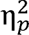 = 0.71, Fig. 4). Accordingly, the velocity that elicited peak power differed with age (*p* < 0.001, 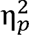 = 0.39) and was faster in the young (355 ± 42 °/s) compared to the old (289 ± 46 °/s; *p* < 0.001, *d* = 0.67) and very old (238 ± 34 °/s; *p* < 0.001, *d* = 1.27) and faster in the old than the very old (*p* < 0.001, *d* = 0.44). Men produced greater absolute power outputs compared to women irrespective of age (*p* < 0.001, 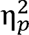 = 0.38) with a larger sex difference at higher velocities (velocity x sex effect, *p* < 0.001, 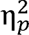 = 0.37) and younger age (age x sex effect, *p* < 0.001, 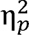 = 0.08, Fig. 4). Peak power was also achieved at a faster velocity in men (309 ± 54 °/s) than women irrespective of age (288 ± 57 °/s; *p* = 0.025; 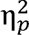 = 0.03).

**Fig. 4.**
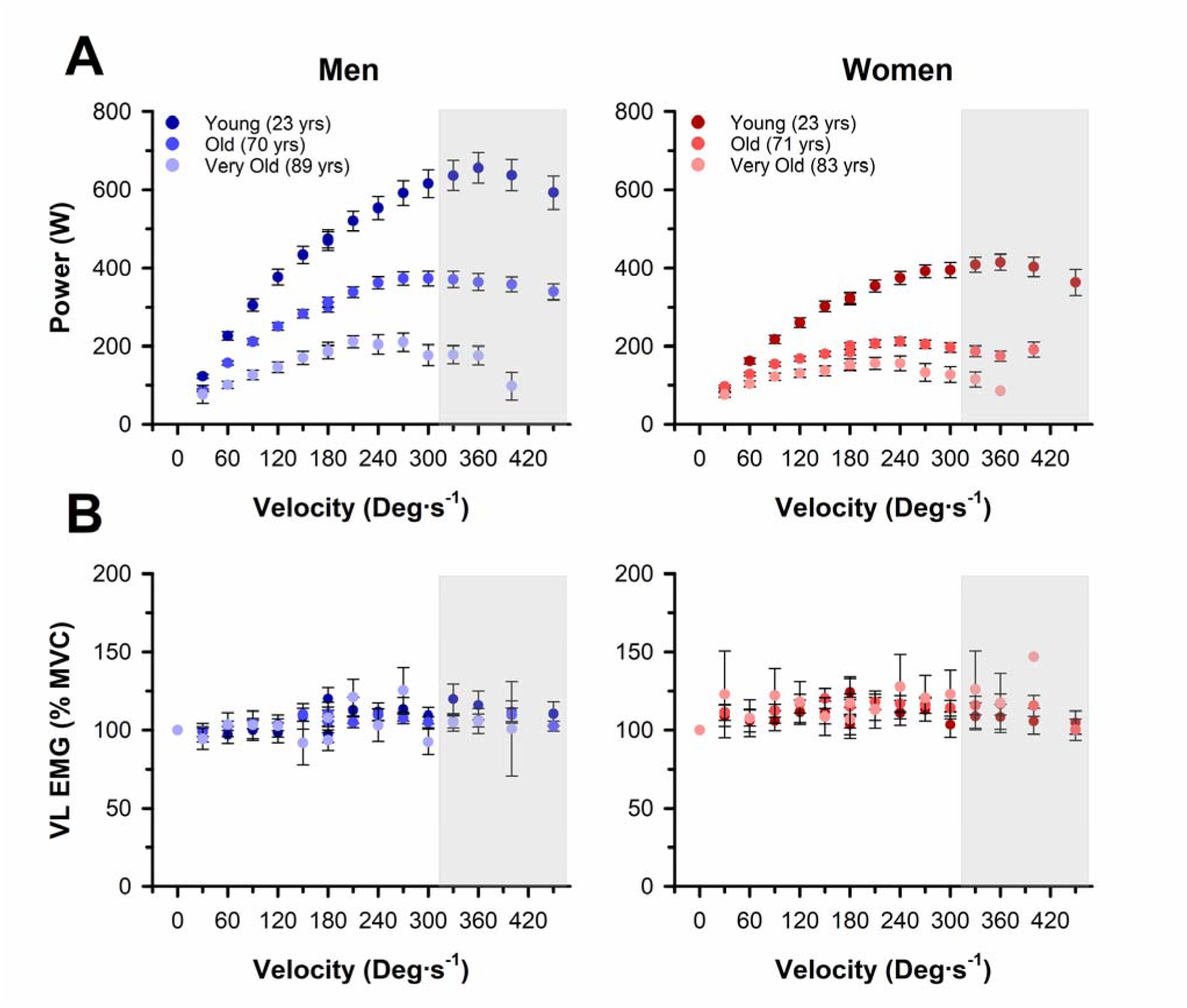
Power output and EMG amplitudes from the vastus lateralis across a range of isokinetic velocities. (A) Young adults produced more power than old and very old adults (*p* < 0.001) with a greater age difference at faster contraction velocities (velocity x age effect, *p* < 0.001). Men produced more power than women (*p* < 0.001) with a greater sex difference at faster contraction velocities (velocity x sex effect, *p* < 0.001). (B) Neural activation, assessed as the surface RMS EMG amplitude of the VL (%MVC), did not differ with age (*p* = 0.631) but was greater in women than men when all ages and velocities were combined (*p* < 0.001). Gray boxes indicate velocities with <100% completion. Values are the means ± SEM.

#### Surface EMG

During the dynamic contractions, and irrespective of the velocity of the contraction, surface RMS EMG amplitude (%MVC) did not differ with age in the VL (*p* = 0.631, 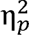 = -0.01, Fig. 4) or VM (*p* = 0.135, 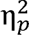 = 0.00) but differed in the rectus femoris with age (*p* < 0.001, 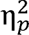 = 0.19). RMS EMG (% MVC) of the rectus femoris was greater in the old than the young (*p* < 0.001, *d* = 0.46) and greater in the very old than the young (*p* < 0.001, d = 1.04) and old (*p* < 0.001, *d* = 0.44). RMS EMG amplitude (%MVC) was greater in women than men for the VL (*p* < 0.001, 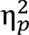 = 0.13), VM (*p* = 0.035, 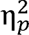 = 0.03), and RF (*p* = 0.003, 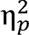 = 0.06), and the sex differences were also greater with increasing age in the VL (age x sex interaction, *p* = 0.020, 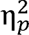 = -0.01, Fig. 4), VM (*p* = 0.003, 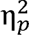 = 0.01), and RF (*p* < 0.001, 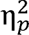 = 0.05). Irrespective of age or sex, there was no effect of the velocity of contraction for the RMS EMG amplitude of the VL (*p* = 0.313, 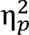 = 0.08), VM (*p* = 0.806, 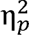 = 0.05), or RF (*p* = 0.909, 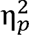 = 0.04).

### Peak Power Output

#### Absolute Peak Power Output

Absolute peak power outputs are reported in Table 2 and displayed in Figure 5. Peak power output for men decreased with age (*p* < 0.001, 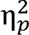 = 0.49) and was 42% and 67% lower in the old (*p* < 0.001, *d* = 1.56) and very old (*p* < 0.001, *d* = 2.49) compared with the young and 41% lower in the very old compared with the old (*p* < 0.001, *d* = 1.12). For women, peak power output decreased with age (*p* < 0.001, 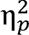 = 0.56) and was 49% and 60% lower in the old (*p* < 0.001, *d* = 2.26) and very old (*p* < 0.001, *d* = 2.88) compared with the young and 23% lower in the very old compared with the old (*p* = 0.016, *d* = 0.74). This corresponded with an age-related loss in peak power output that was ∼6.5 W·year^-1^ for men (*R*^2^ = 0.62, *p* < 0.001), which was a greater rate of decline (*p* = 0.002, 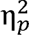 = 0.07) than the ∼4.2 W·year^-1^ for women (*R*^2^ = 0.77, *p* < 0.001). In contrast, when we expressed the peak power outputs relative to the mean of the young men and women, the relative rate of decline did not differ in the men (∼1.0 %·year^-1^) compared with women (∼1.0 %·year^-1^, *p* = 0.705, 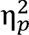 = 0.00). Excluding the young adults from the regression analyses resulted in an age-related loss in peak power output of ∼10.7 W·year^-1^ for men (*R*^2^ = 0.43, *p* < 0.001), which was a greater rate of decline (*p* = 0.002, 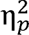 = 0.10) than the ∼3.8 W·year^-1^ for women (*R*^2^ = 0.22, *p* < 0.001). In contrast, the relative rate of decline in peak power output in old and very old did not differ between the men (∼1.5 %·year^-1^) and women (∼0.9 %·year^-1^, *p* = 0.070, 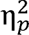 = 0.04).

**Fig. 5.**
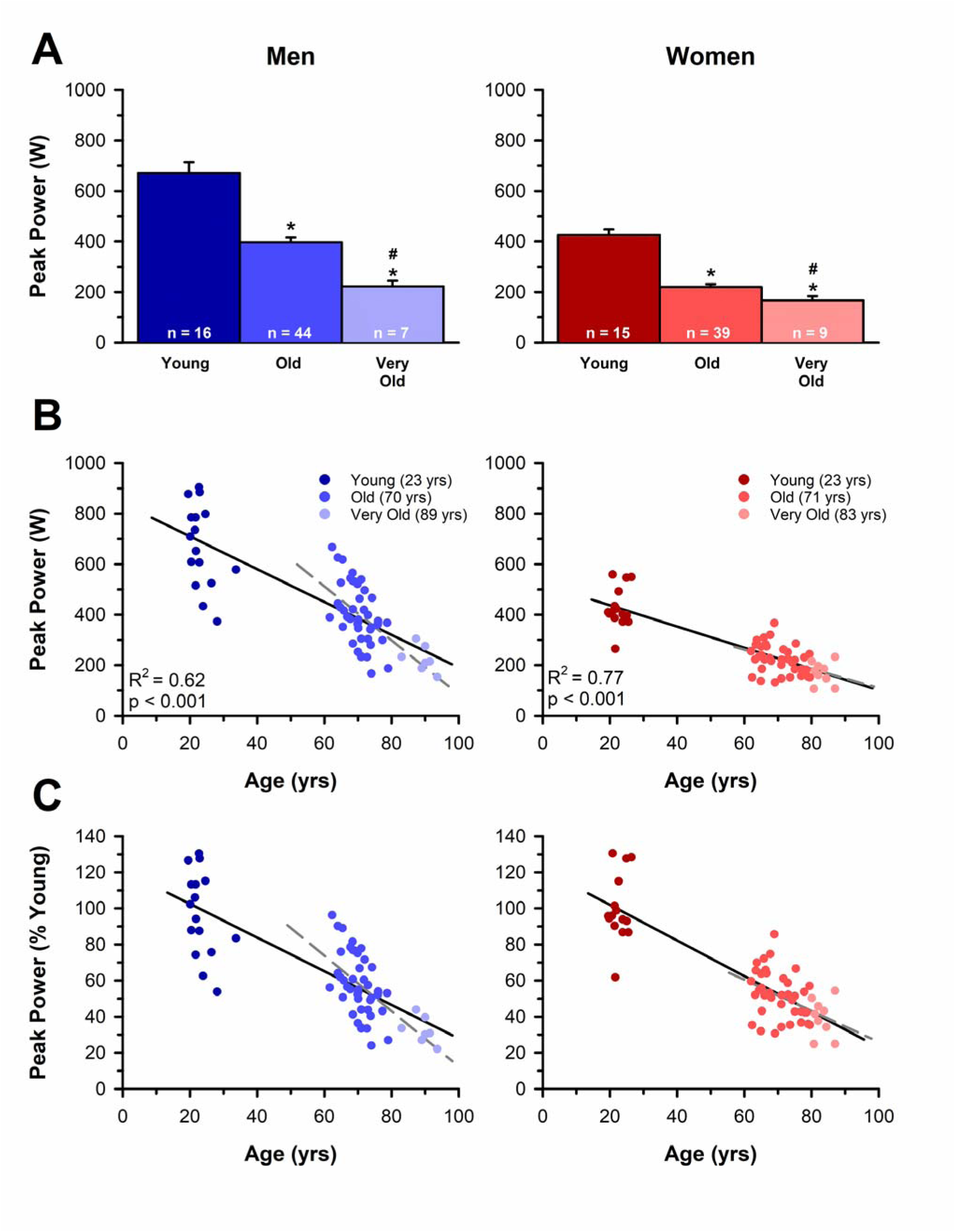
Decrements in absolute peak power with age. (A) Absolute peak mechanical power outputs for young, old, and very old men and women. (B) The age-related loss in peak power output was ∼6.5 W· year^-1^ for men, which was a greater rate of decline (*p* = 0.002) than the ∼4.2 W· year^-1^ for women. Excluding the young adults from the regression analyses resulted in an age-related loss in peak power output of ∼10.7 W· year^-1^ for men (*R*^2^ = 0.43, *p* < 0.001), which remained a greater rate of decline (*p* = 0.002) than the ∼3.8 W·year^-1^ for women (*R*^2^ = 0.22, *p* < 0.001). (C) The reduction in peak power relative to the mean peak power of the young, sex-matched adults did not differ (*p* = 0.705) in the men (∼1.0 %·year^-1^) compared with women (∼1.0 %·year^-1^). When the young adults were excluded from the regression analyses, the relative rate of power loss remained not different (*p* = 0.070) between the men (∼1.5 %·year^-1^) and women (∼0.9 %·year^-1^). Values in (A) are the means ± SEM. Black solid lines and gray dashed lines in (B) and (C) are the least squares regression lines for all three age cohorts and excluding the young adults, respectively.

#### Mass-Specific Peak Power Output

Mass-specific power outputs are reported in Table 2 and displayed in Figure 6. Mass-specific peak power output for men decreased with age (*p* < 0.001, 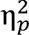 = 0.53) and was 35% and 53% lower the old (*p* < 0.001, *d* = 1.92) and very old (*p* < 0.001, *d* = 2.49) compared with the young and 28% lower in the very old compared with the old (*p* = 0.004, *d* = 0.86).For women, mass-specific peak power output also decreased with age (*p* < 0.001, 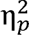 = 0.60) and was 38% and 55% lower in the old (*p* < 0.001, *d* = 2.33) and very old (*p* < 0.001, *d* = 2.88) compared with the young and 28% lower in the very old compared with the old (*p* = 0.004, *d* = 0.90). This corresponded with an age-related loss in mass-specific peak power output that was ∼0.74 W·kg^-1^·year^-1^ for men (*R* = 0.64, *p* < 0.001) and ∼0.67 W·kg ·year for women (*R* = 0.68, *p* < 0.001), which did not differ between the sexes (*p* = 0.452, 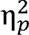 = 0.01, Fig. 6). Similarly, when we expressed the mass-specific peak power outputs relative to the mean of the young men and women, the relative rate of decline did not differ in the men (∼0.8 %·year^-1^) compared with women (∼0.8 %·year^-1^, *p* = 0.676, 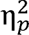 = 0.00). Excluding the young from the regression analyses resulted in an age-related loss in mass-specific peak power output of ∼1.09 W·kg^-1^·year^-1^ for men (*R*^2^ = 0.33, *p* < 0.001) and ∼0.91 ·kg^-1^·year^-1^ for women (*R*^2^ = 0.27, *p* < 0.001), which did not differ between the sexes (*p* = 0.579, 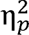 = 0.00). Similarly, the relative rate of decline in mass-specific peak power output in old and very old did not differ in the men (∼1.2 %·year^-1^) compared with women (∼1.1 %·year^-1^, *p* = 0.939, 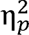 = 0.00).

**Fig. 6.**
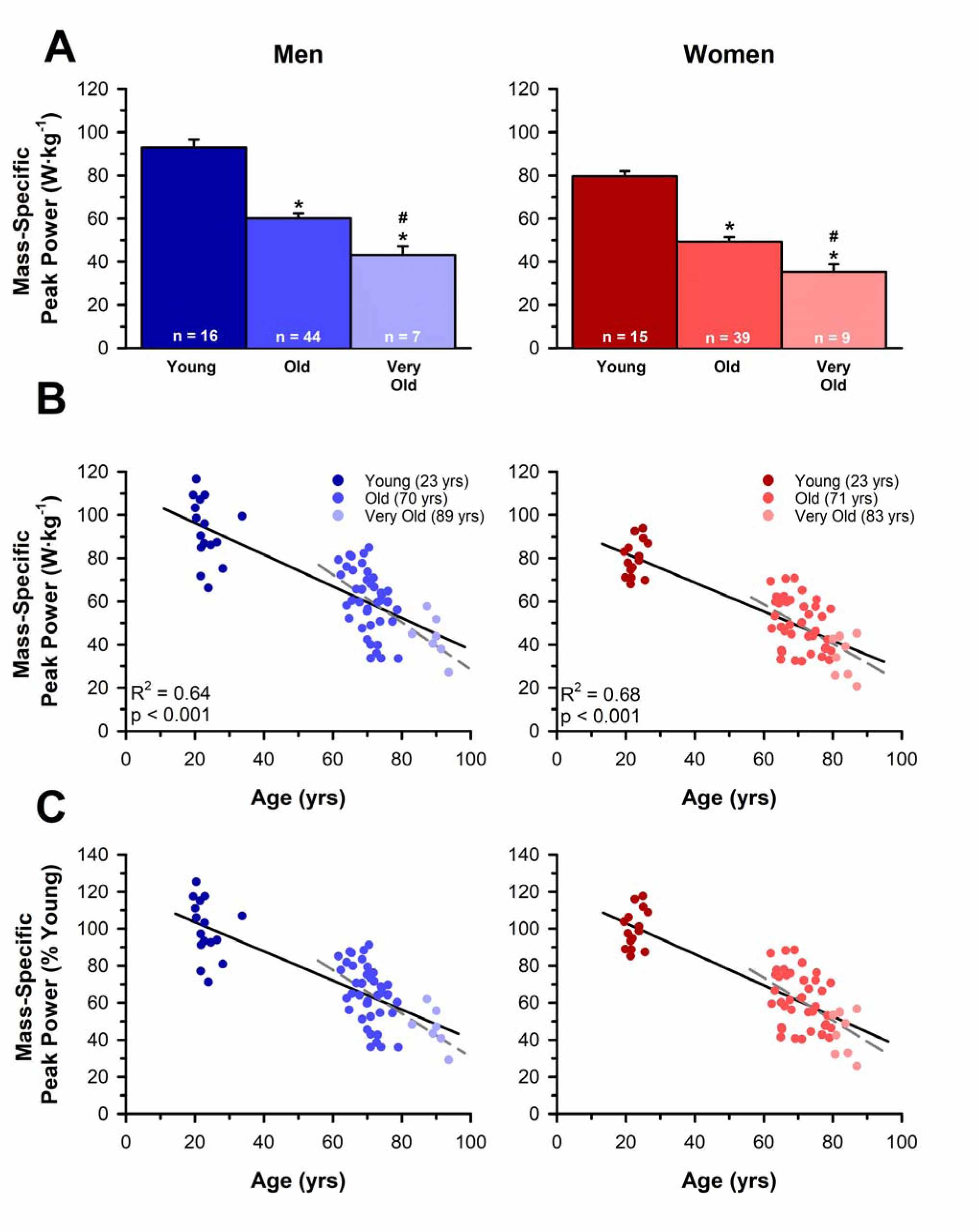
Decrements in mass-specific peak power with age. (A) Mass-specific peak power outputs for young, old, and very old men and women. (B) The age-related loss in mass-specific peak power output was ∼0.74 W·kg^-1^·year^-1^ for men and ∼0.67 W·kg^-1^·year^-1^ for women, which did not differ between the sexes (*p* = 0.452). Excluding the young adults from the regression analyses resulted in a loss in mass-specific peak power output of ∼1.09 W·kg^-1^·year^-1^ for men (*R*^2^ = 0.33, *p* < 0.001) and ∼0.91 ·kg^-1^·year^-1^ for women (*R*^2^= 0.27, *p* < 0.001), which also did not differ between the sexes (*p* = 0.579). (C) The reduction in peak power relative to the mean peak power of the young, sex-matched adults did not differ (*p* = 0.676) in the men (∼0.8 %·year^-1^) compared with women (∼0.8 %·year^-1^). When the young adults were excluded from the regression analyses, the relative rates of mass-specific peak power loss remained not different (*p* = 0.939) between the men (∼1.2 %·year^-1^) and women (∼1.1 %·year^-1^). Values in (A) are the means ± SEM. Black solid lines and gray dashed lines in (B) and (C) are the least squares regression lines for all three age cohorts and excluding the young adults, respectively.

### Neuromuscular Stimulation Parameters and Associations with Peak Power

#### Involuntary Contractile Properties

Neuromuscular stimulation measures are reported in Table 2. Due to the large number of involuntary contractile property measurements, we only report the three variables with the strongest associations with peak power in the text. Potentiated twitch amplitude differed with age (*p* < 0.001, 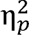 = 0.16) and was greater in the young than the old (*p* = 0.002, *d* = 0.64) and very old adults (*p* < 0.001, *d* = 1.60), greater in the old than the very old adults (*p* = 0.005, *d* = 0.61), and greater in men than women (*p* < 0.001, 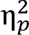 = 0.39). Rate of twitch torque development differed with age (*p* < 0.001, 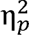 = 0.20) and was greater in the young than the old (*p* < 0.001, *d* = 0.68) and very old adults (*p* < 0.001, *d* = 1.96), greater in the old than very old adults (*p* = 0.001, *d* = 0.70), and greater in men than women (*p* < 0.001, 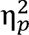 = 0.37). Peak rate of torque relaxation following TMS differed with age (*p* < 0.001, 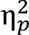 = 0.12) and was faster in the young than the old (*p* < 0.001, d = 0.69) and very old adults (*p* < 0.001, *d* = 1.37) and faster in men than women (*p* < 0.001, 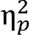 = 0.32), but did not differ between the old and very old adults (*p* = 0.242, *d* = 0.25).

Pearson correlation coefficients between peak power and the neuromuscular stimulation parameters are reported in Table 3 for men and women and in Table 4 for young, old, and very old adults. For men, muscular factors, including lean thigh tissue mass (*R*^2^ = 0.53, *p* < 0.001) and rate of twitch torque development (*R*^2^ = 0.69, *p* < 0.001) were strongly associated with peak power (Fig. 7). Similarly, lean thigh tissue mass (*R*^2^ = 0.34, *p* < 0.001) and rate of twitch torque development (*R*^2^ = 0.57, *p* < 0.001) were strongly associated with peak power in women (Fig. 7). To account for the influence of muscle size on both peak power and the involuntary electrically evoked contractile measurements, associations with mass-specific power were also assessed. After normalizing peak power to the lean thigh tissue mass, the association with the rate of twitch torque development remained strong in men (*R*^2^ = 0.38, *p* < 0.001) and women (*R*^2^ = 0.38, *p* <.001, Fig. 7).

**Fig. 7.**
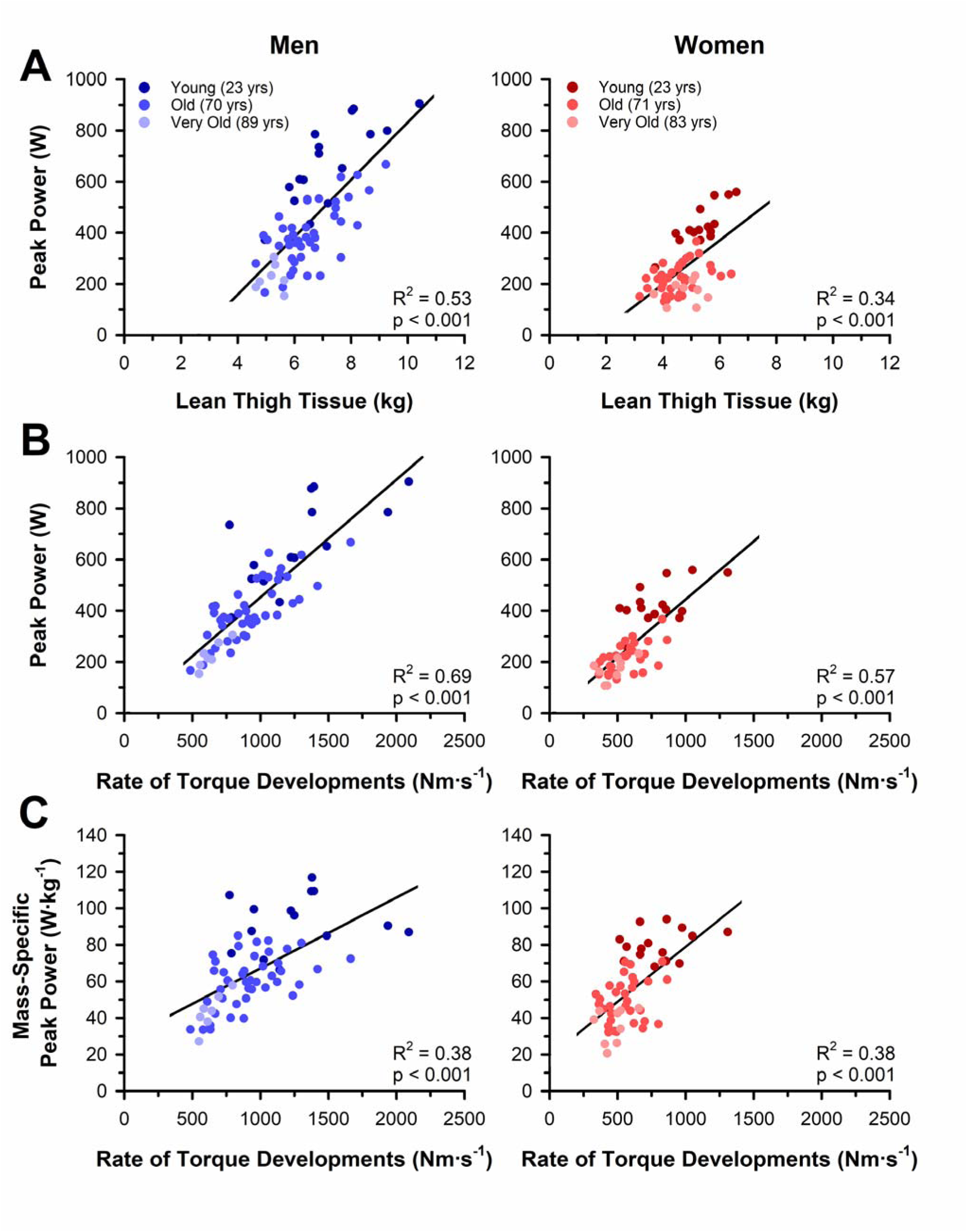
Associations between peak power and muscular variables in men and women. (A) Lean thigh tissue mass was closely associated with absolute peak power in men and women. (B) Rates of twitch torque development were also closely associated with absolute peak power in men and women. (C) The strength of the association between peak power and the rate of torque development remained strong for men and women when peak power was normalized to lean thigh tissue mass.

**Table 3:** Pearson correlation coefficients between peak power and neuromuscular performance measures in men and women.

| Variable | Men (65) |  | Women (57) |  |
| --- | --- | --- | --- | --- |
| | Absolute<br>Peak Power (W) | Mass-Specific<br>Peak Power ($\text{W} \cdot \text{kg}^{-1}$) | Absolute<br>Peak Power (W) | Mass-Specific<br>Peak Power ($\text{W} \cdot \text{kg}^{-1}$) |
| Thigh Lean Tissue Mass | 0.725** | ----- | 0.583** | ----- |
| Twitch Torque - $Q_{\text{tw}}$ | 0.789** | 0.533** | 0.759** | 0.585** |
| Rate of Torque Dev. | 0.827** | 0.613** | 0.758** | 0.618** |
| Norm. Rate of Torque Dev. | 0.266* | 0.358** | 0.195 | 0.218 |
| ½ Relaxation Time | -0.379** | -0.287* | -0.17 | -0.15 |
| Estimated Resting Twitch | 0.477** | 0.344** | 0.501** | 0.288* |
| Peak Relaxation Rate | -0.723** | -0.527** | -0.621** | -0.514** |
| Norm. Peak Relaxation Rate | -0.023 | 0.028 | 0.123 | 0.107 |
| Voluntary Activation (SIT/ERT) | 0.164 | 0.258* | 0.337* | 0.409** |
| Voluntary Activation (SIT/$Q_{\text{tw}}$) | 0.232 | 0.301* | 0.287* | 0.376** |
| VL RMS EMG at Peak Power | 0.113 | 0.119 | 0.02 | 0.039 |
| VM RMS EMG at Peak Power | 0.076 | 0.138 | 0.02 | 0.084 |
| RF RMS EMG at Peak Power | -0.091 | -0.162 | -0.235 | -0.146 |

The sample sizes ( $n$ ) for each cohort are reported in parentheses. Removing the voluntary activation for the very old woman outlier resulted in minimal changes to the association between voluntary activation and absolute peak power (*eq. 1* (SIT/ERT): $r = 0.303$ , $p = 0.021$ ; *eq. 2* (SIT/ $Q_{\text{tw}}$ ): $r = 0.236$ , $p = 0.083$ ) and mass-specific peak power (*eq. 1* (SIT/ERT): $r = 0.385$ , $p = 0.004$ ; *eq. 2* (SIT/ $Q_{\text{tw}}$ ): $r = 0.353$ , $p = 0.008$ ) in women. \* and \*\* indicate a significant correlation at $p \leq 0.05$ and $p \leq 0.01$ , respectively.

**Table 4:**
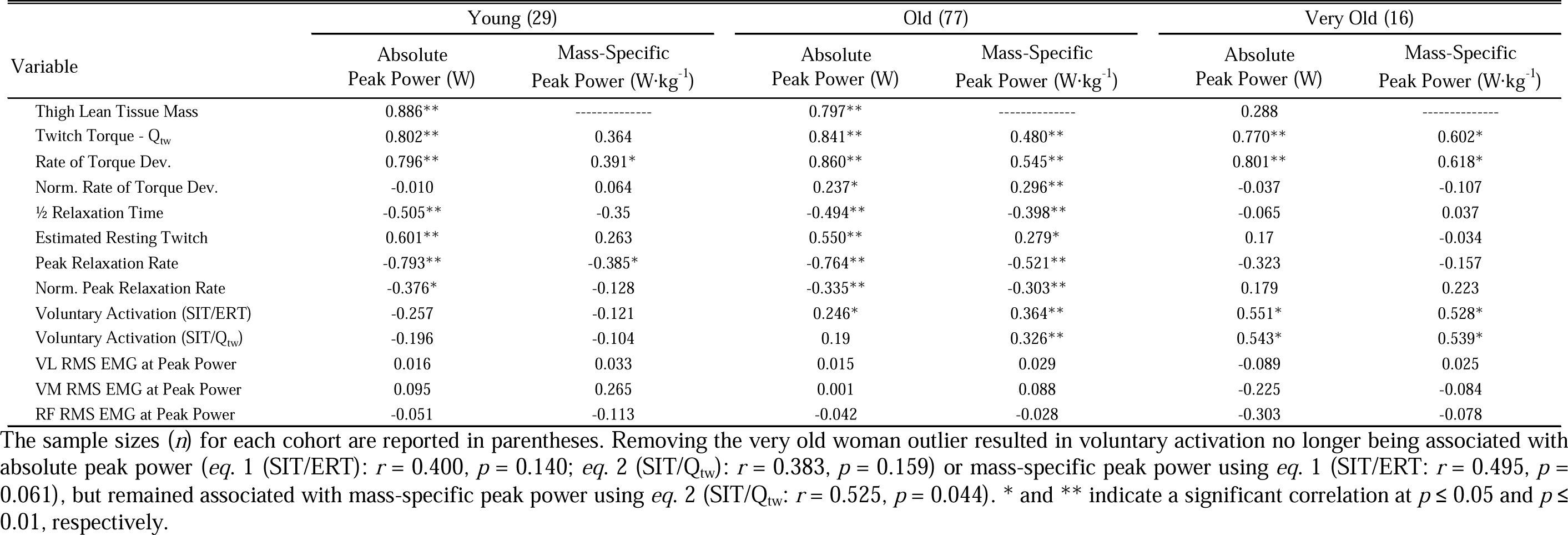
Pearson correlation coefficients between peak power and neuromuscular performance measures in young, old, and very old adults.

| Variable | Young (29) |  | Old (77) |  | Very Old (16) |  |
| --- | --- | --- | --- | --- | --- | --- |
|  | Absolute Peak Power (W) | Mass-Specific Peak Power (W·kg<sup>-1</sup>) | Absolute Peak Power (W) | Mass-Specific Peak Power (W·kg<sup>-1</sup>) | Absolute Peak Power (W) | Mass-Specific Peak Power (W·kg<sup>-1</sup>) |
| Thigh Lean Tissue Mass | 0.886** | ----- | 0.797** | ----- | 0.288 | ----- |
| Twitch Torque - Q<sub>tw</sub> | 0.802** | 0.364 | 0.841** | 0.480** | 0.770** | 0.602* |
| Rate of Torque Dev. | 0.796** | 0.391* | 0.860** | 0.545** | 0.801** | 0.618* |
| Norm. Rate of Torque Dev. | -0.010 | 0.064 | 0.237* | 0.296** | -0.037 | -0.107 |
| ½ Relaxation Time | -0.505** | -0.35 | -0.494** | -0.398** | -0.065 | 0.037 |
| Estimated Resting Twitch | 0.601** | 0.263 | 0.550** | 0.279* | 0.17 | -0.034 |
| Peak Relaxation Rate | -0.793** | -0.385* | -0.764** | -0.521** | -0.323 | -0.157 |
| Norm. Peak Relaxation Rate | -0.376* | -0.128 | -0.335** | -0.303** | 0.179 | 0.223 |
| Voluntary Activation (SIT/ERT) | -0.257 | -0.121 | 0.246* | 0.364** | 0.551* | 0.528* |
| Voluntary Activation (SIT/Q<sub>tw</sub>) | -0.196 | -0.104 | 0.19 | 0.326** | 0.543* | 0.539* |
| VL RMS EMG at Peak Power | 0.016 | 0.033 | 0.015 | 0.029 | -0.089 | 0.025 |
| VM RMS EMG at Peak Power | 0.095 | 0.265 | 0.001 | 0.088 | -0.225 | -0.084 |
| RF RMS EMG at Peak Power | -0.051 | -0.113 | -0.042 | -0.028 | -0.303 | -0.078 |

The sample sizes (*n*) for each cohort are reported in parentheses. Removing the very old woman outlier resulted in voluntary activation no longer being associated with absolute peak power (*eq. 1* (SIT/ERT): $r = 0.400$ , $p = 0.140$ ; *eq. 2* (SIT/Q<sub>tw</sub>): $r = 0.383$ , $p = 0.159$ ) or mass-specific peak power using *eq. 1* (SIT/ERT: $r = 0.495$ , $p = 0.061$ ), but remained associated with mass-specific peak power using *eq. 2* (SIT/Q<sub>tw</sub>: $r = 0.525$ , $p = 0.044$ ). \* and \*\* indicate a significant correlation at $p \leq 0.05$ and $p \leq 0.01$ , respectively.

For young adults, muscular factors, including lean thigh tissue mass (*R*^2^ = 0.78, *p* < 0.001) and rate of twitch torque development (*R*^2^ = 0.63, *p* < 0.001) were strongly associated with peak power. Similarly, lean thigh tissue mass (*R*^2^ = 0.64, *p* < 0.001) and rate of twitch torque development (*R*^2^ = 0.74, *p* < 0.001) were strongly associated with peak power in old adults. However, the thigh lean tissue mass of very old adults was not associated with peak power (*R*^2^ = 0.08, *p* = 0.279), but the rate of twitch torque development was strongly associated with peak power (*R*^2^ = 0.64, *p* = 0.004). After normalizing peak power to lean thigh tissue mass, the associations with the rate of twitch torque development remained in the young (*R*^2^ = 0.15, *p* = 0.036), old (*R*^2^ = 0.30, *p* < 0.001), and very old (*R*^2^ = 0.38, *p* = 0.011).

#### Voluntary Activation

Voluntary activation did not differ with age or sex when calculated by *eq. 1* that uses the estimated resting twitch (age: *p* = 0.163, 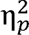 = 0.01; sex: *p* = 0.374, 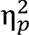 = 0.00) or *eq. 2* that uses the potentiated resting twitch (age: *p* = 0.337, 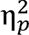 = 0.00; sex: *p* = 0.695, 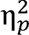 = 0.01).

Voluntary activation was not associated with peak power in men (*eq. 1*: *R*^2^ = 0.03, *p* = 0.191, Fig. 8; *eq. 2*: *R*^2^ = 0.05, *p* = 0.063) but was weakly associated with peak power in women (*eq. 1*: *R*^2^ = 0.12, *p* = 0.012, Fig. 8; *eq. 2*: *R*^2^ = 0.08, *p* = 0.032). Importantly, removing the very old woman outlier resulted in minimal changes to the association between voluntary activation and peak power in women (*eq. 1*: *R*^2^ = 0.10, *p* = 0.021; *eq. 1*: *R*^2^ = 0.06, *p* = 0.083). Voluntary activation was also not associated with peak power in the young adults (*eq. 1*: *R*^2^ = 0.07, *p* = 0.179; *eq. 2*: *R*^2^ = 0.04, *p* = 0.308) but was associated with peak power in the very old adults (*eq. 1*: *R*^2^ = 0.31, *p* = 0.027; *eq. 2*: *R*^2^ = 0.30, *p* = 0.030) and the old adults when using *eq. 1* (*R*^2^ = 0.07, *p* = 0.033) but not *eq. 2* (*R*^2^ = 0.04, *p* = 0.100). When the very old woman outlier was removed from the analyses, voluntary activation was no longer associated with peak power in the very old adults (*eq. 1*: *R*^2^ = 0.16, *p* = 0.140; *eq. 2*: *R*^2^ = 0.15, *p* = 0.159).

**Fig. 8.**
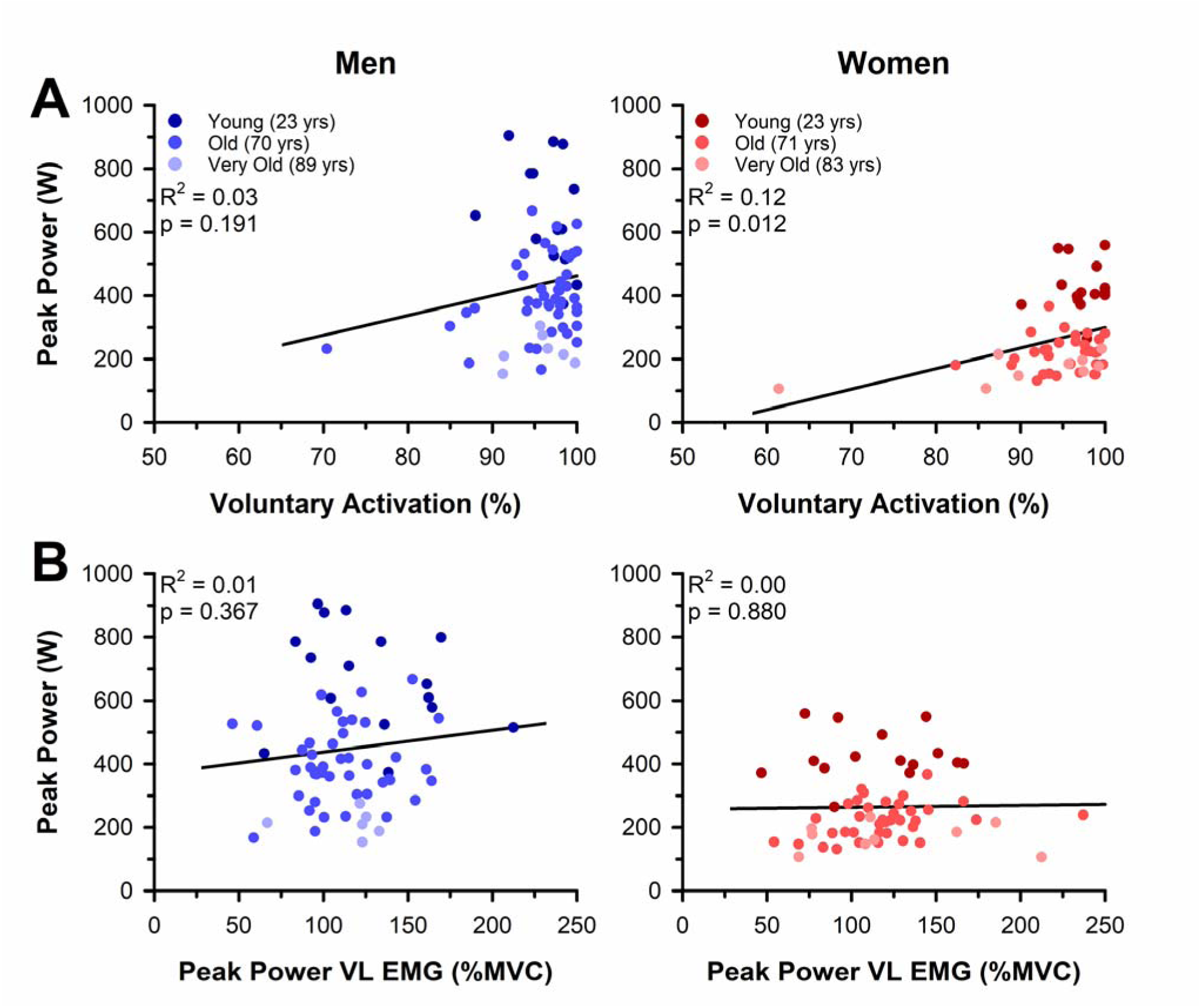
Associations between peak power and neural activation in men and women. (A) Voluntary activation assessed with TMS delivered to the motor cortex during the isometric MVCs was weakly associated with absolute peak power in women but not in men. Removing the very old woman outlier resulted in minimal changes to the strength of the association between voluntary activation and peak power in women (*R*^2^ = 0.10, *p* = 0.021). (B) VL RMS EMG amplitude (%MVC) eliciting peak power was not associated with absolute peak power for men (*R*^2^ = 0.01, *p* = 0.367) or women (*R*^2^ = 0.00, *p* = 0.880).

#### Surface EMG Eliciting Peak Power

VL RMS EMG amplitude (%MVC) eliciting peak power did not differ with age (*p* = 0.726, 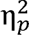 = 0.01) or sex (*p* = 0.627, 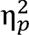 = 0.01). Similarly, the VL RMS EMG amplitude (%MVC) eliciting peak power was not associated with peak power in men (*R*^2^ = 0.01, *p* = 0.367) or women (*R*^2^ = 0.00, *p* = 0.880, Fig. 8), or in the young (*R*^2^ = 0.00, *p* = 0.931), old (*R*^2^ = 0.00, *p* = 0.897), or very old (*R*^2^ = 0.01, *p* = 0.754).

### MEPs and Silent Period

Peak-to-peak MEPs (%M_max_) of the VL did not differ with age (*p* = 0.056, 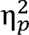 = 0.03) or sex (*p* = 0.085, 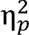 = 0.02) and were not associated with peak power in men (*R*^2^ = 0.00, *p* = 0.625) or women (*R*^2^ = 0.00, *p* = 0.992) nor in young (*R*^2^ = 0.05, *p* = 0.231), old (*R*^2^ = 0.01, *p* = 0.511), or very old (*R*^2^ = 0.14, *p* = 0.147). The cortical silent period of the VL following the TMS however, differed with age (*p* < 0.001, 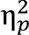 = 0.22) and was longer in the young than the old (*p* < 0.001, *d* = 1.22) and very old (*p* = 0.006, *d* = 0.90) but did not differ between men and women (*p* = 0.115, 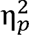 = 0.01) or between old and very old (*p* = 0.344, *d* = 0.20). Accordingly, the cortical silent period of the VL was associated with peak power in the men (*R*^2^ = 0.14, *p* = 0.002) and women (*R*^2^ = 0.19, *p* < 0.001) but was not associated with peak power in the young (*R*^2^ = 0.04, *p* = 0.330), old (*R*^2^ = 0.03, *p* = 0.135), or very old (*R*^2^ = 0.03, *p* = 0.505).

### Physical Activity

Physical activity (steps·day^-1^) differed with age (*p* < 0.001, 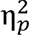 = 0.21) and was greater in the young than the very old (*p* < 0.001, *d* = 1.87) and greater in the old than the very old (*p* < 0.001, *d* = 1.03) but did not differ between the young and old (*p* = 0.061, *d* = 0.40) or between men and women (*p* = 0.557, 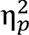 = 0.01). Physical activity (steps·day^-1^) was associated with peak power in women (*R*^2^ = 0.12, *p* = 0.012) but not in men (*R*^2^ = 0.00, *p* = 0.698), young (*R*^2^ = 0.06, *p* = 0.250), old (*R*^2^ = 0.00, *p* = 0.751), or very old adults (*R*^2^ = 0.24, *p* = 0.057).

### Functional Performance

Functional performance test scores for the old and very old adults are reported in Table 5, and the associations with peak power are displayed in Supplemental Figure 1. Berg Balance Test score was higher in the old than the very old adults (*p* < 0.001, 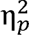 = 0.26) but did not differ between men and women (*p* = 0.983, 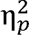 = -0.01). Six-minute walk test distance was greater in the old than the very old adults (*p* < 0.001, 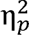 = 0.14) and greater in men than women (*p* = 0.004, 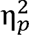 = 0.08). Stair climb time was faster in the old than the very old adults (*p* < 0.001, 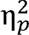 = 0.20) but did not differ between men and women (*p* = 0.056, 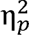 = 0.03). Sit-to-stand test time (5x) was faster in the old than the very old adults (*p* < 0.001, 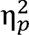 = 0.13) but did not differ between men and women (*p* = 0.190, 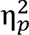 = 0.01). Sit-to-stand test time (10x) was also faster in the old than the very old adults (*p* < 0.001, 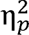 = 0.12) but did not differ between men and women (*p* = 0.176, 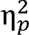 = 0.01). Absolute peak power of the knee extensors was associated with the Berg Balance Test (*R*^2^ = 0.08, *p* = 0.007), six-minute walk distance (*R*^2^ = 0.28, *p* < 0.001), stair climb time (*R*^2^ = 0.17, *p* < 0.001), and sit-to-stand scores (5x: *R*^2^ = 0.10, *p* = 0.003; 10x: *R*^2^ = 0.08, *p* = 0.007).

**Table 5:** Functional performance scores from the old and very old men and women.

| Variable | Units | Old (60-79 yrs) | | Very Old ($\geq 80$ yrs) | |
| --- | --- | --- | --- | --- | --- |
|  |  | Men (41) | Women (35) | Men (7) | Women (9) |
| Berg Balance Test* | Score | 55.0 ± 1.6 (40) | 54.7 ± 2.4 (34) | 49.4 ± 6.8 | 49.6 ± 4.7 (8) |
| 6-minute walk test distance*† | m | 597.9 ± 60.7 | 546.2 ± 66.4 | 442.3 ± 129.6 | 459.0 ± 120.6 |
| Stair climb time* | s | 5.0 ± 1.4 (40) | 5.4 ± 1.7 | 6.4 ± 1.7 | 8.2 ± 1.5 |
| Sit-to-stand time (5x)* | s | 9.2 ± 1.8 (40) | 9.6 ± 2.5 (34) | 12.0 ± 5.2 (6) | 13.7 ± 2.7 |
| Sit-to-stand time (10x)* | s | 19.0 ± 3.7 | 19.7 ± 5.4 (34) | 23.2 ± 8.0 (6) | 28.6 ± 6.3 |

The sample sizes (*n*) for each cohort and certain variables are reported in parentheses. One very old man failed the sit-to-stand test (5x and 10x). Symbols next to the variable name indicate a significant effect of \* age, † sex, or an ‡ age x sex interaction at $p \leq 0.05$ . Values are reported as means ± SD

## DISCUSSION

This study determined the loss in power output of the knee extensor muscles with aging across a wide range of velocities and identified the neural and muscular contributions to peak power loss in old and very old men and women. As expected, we observed that aging was associated with a progressive loss in isometric strength and power output, with the greatest impairments in power occurring at the faster contraction velocities. The rate of decline in absolute maximal isometric strength and power output with age was greater in men than women; however, the relative rate of decline did not differ between the sexes. We provide evidence that voluntary activation is at or near maximal for the young, old, and very old adults (Table 2, Fig. 4) but may play a minor role for the age-related peak power loss in older women (Table 4, Fig. 8). Other neural indices including the cortical silent period and MEP also did not explain the loss of muscle power with age. The mechanisms for the loss in power with age appear largely similar between men and women, with thigh lean tissue mass and involuntary electrically evoked contractile properties emerging as the most closely associated variables with peak power (Tables 3 & 4, Fig. 7). Importantly, when peak power was normalized to thigh lean tissue mass, the involuntary contractile properties remained closely associated with power. These data suggest that the age-related loss in muscle mass and other factors altering the contractile properties of the muscle are the primary mechanisms for the loss in power of old and very old men and women.

### Rates of Peak Power Loss with Aging in Men Compared to Women

Our data shows that the rate of absolute peak power loss with aging is greater in men (∼6.5 W·year^-1^) than women (∼4.2 W·year^-1^; Fig. 5). This finding is consistent with previous cross-sectional (2-6, 50) and longitudinal studies (3, 4, 51) that found a greater rate of absolute strength and power loss in men. However, the rates of power loss expressed relative to the amount of thigh lean tissue mass did not differ between the men (∼0.74 W·kg^-1^·year^-1^) and women (∼0.67 W·kg^-1^·year^-1^; Fig. 6), suggesting that the sex differences in absolute power loss with aging are attributed, in large part, to the differences in muscle mass between men and women that persist across the lifespan (2, 6, 51-55). This finding is consistent with previous whole-muscle (2, 56) and single-fiber (56) studies that also found changes in muscle size to account for a majority of the sex differences in the loss of power output with age. In addition, when the power outputs were expressed as a percentage of the respective means from the young adults, the yearly loss in power (∼1.0%) and mass-specific power (∼0.8%) did not differ between the sexes. Several other studies have also observed no sex difference in the relative reduction in power with aging (2-4), suggesting that the greater absolute rate of age-related power loss in men is likely attributed to the greater power outputs and muscle mass present in young men compared with women prior to aging.

An important limitation in the present study is the rates of power loss with aging are estimated based on cross-sectional data spanning several decades where power output has been shown to be relatively stable (2-4). Thus, our data most likely underestimates the rate of power loss that occurs after the age of ∼40-50 years old. Indeed, other studies have found an annual power loss of ∼2-4% in older adults (2, 3, 5), which is much greater than the ∼1% observed in the present study. To address this limitation, we excluded the young adults from our regression analyses and found that the estimated power loss increased in older men (∼10.7 W·year^-1^ and ∼1.5%·year^-1^) but was unchanged, or even attenuated, in older women (∼3.8 W·year^-1^ and ∼0.9 %·year^-1^). This apparent sex difference in power loss during advanced age (≥60 years) may be due to an unknown mechanism(s) that prevents power loss in older women, or the more likely explanation of limitations with cross-sectional study designs, such as the survivorship bias or “floor effect” that has been proposed in previous studies observing a preservation of power in very old women (2, 5). Future studies aimed at studying older women with more advanced age and low physical function are needed to distinguish between physiological differences and sampling bias.

In addition to the attenuated rate of power loss in older women, our findings observed a steady rate of power loss among old and very old men, in contrast to the accelerated rate of power loss in very old adults reported in some previous studies (2, 4, 5). The explanation for the discrepancies between our study and others is unknown, but one likely possibility is the same survivorship bias and floor effect that was proposed for the attenuated loss of power in very old women, along with the health inclusion criteria for this study. Notably, the majority of the old and very old men and women in this study scored above clinical cutoffs on several common tests of physical function (e.g., Berg Balance, 97.8% scored ≥ 45 (57); 6MWT, 93.5% ≥ 400m (58, 59); 5x sit-to-stand, 95.6% ≤ 15s (60)). Therefore, the findings of this study and others using a cross-sectional study design in the laboratory setting likely underestimate the rate of power loss in the general population of older adults, especially in very old adults and potentially older women who have the lowest power outputs.

### Age-Related Loss in Power is Determined Primarily by Mechanisms Within the Muscle for Men and Women

In the present study, we integrated measures of surface EMG with TMS to the motor cortex and electrical stimulation to the femoral nerve to localize the primary sites along the motor pathway contributing to the age-related loss in power in men compared with women. Although we did not observe a difference in the ability of the motor cortex to volitionally activate the knee extensor muscles with age (Table 2), we did observe a significant, albeit weak, association between voluntary activation and peak power output in women that was not present in men (Fig. 8). These findings suggest that a reduced ability of the motor cortex to voluntarily activate the knee extensor muscles may contribute, at least in part, to the age-related loss in power of women. It should be noted, however, that methodological limitations with the interpolated twitch technique to assess voluntary activation preclude its use during the moderate-to high-velocity contractions necessary to obtain peak power. In recognition of this limitation, we measured the RMS EMG amplitude of the rectus femoris, vastus medialis, and vastus lateralis muscles during the dynamic contractions and found that the RMS EMG amplitude either did not differ with age (Fig. 4) or was slightly higher in the old and very old adults across the power-velocity relationship. There were also no associations between the RMS EMG amplitude and peak power of the knee extensors in men or women (Table 3, Fig. 8) or in the old and very old adults (Table 4). These data are in agreement with a majority of the literature from both isometric and slow dynamic contractions that report minimal to no age differences in voluntary activation of several muscle groups when older adults are familiarized to the experimental procedures (7, 24, 25, 61, 62). Thus, interpreted together, these data provide compelling evidence that a reduced ability to maximally activate the knee extensor muscles plays a minimal role in the age-related loss in force and power output for mobile, community-dwelling old and very old adults. These findings do not likely extend to mobility-impaired, frail older adults, however, as there is evidence for greater impairments in neural activation of the knee extensor muscles in this older adult cohort (36, 37, 55, 63).

In contrast to the limited involvement of the nervous system in explaining the loss of power with aging, we observed close associations between peak power and the age-related loss of thigh lean tissue mass and other mechanisms that disrupt contractile function within the knee extensor muscles (Tables 3 and 4). Specifically, thigh lean tissue mass and the rate of torque development from the involuntary electrically evoked twitch explained 53% and 69% of the variance in peak power in men and 34% and 57% of the variance in women (Fig. 7). Although we observed no association between peak power and thigh lean tissue mass in the very old adults, we postulate this was likely due to methodological limitations as DXA has been shown to underestimate the loss of muscle mass with aging (43), particularly in the knee extensors which appear to atrophy at a greater rate than the knee flexors (43, 52, 64). Irrespective of the potential explanation, the amplitude and rate of torque development of the involuntary electrically evoked twitch were strongly associated with peak power in the very old adults, explaining 59% and 64% of the variance, respectively (Table 4). Importantly, these associations remained strong after normalizing peak power to the thigh lean tissue mass in both the old and very old adults (Table 4) and in men and women (Fig. 7, Table 3). These findings suggest that factors intrinsic to the muscle that slow the involuntary twitch properties are a primary mechanism for the loss in peak power in old and very old men and women.

Although the specific mechanisms within the muscle cannot be identified from the electrically evoked twitch, interpreting our findings in conjunction with the prevailing literature can help shed light on the most likely mechanisms for the accelerated loss in power output relative to muscle mass with aging. Several factors in the muscle that may explain this phenomenon include, but are not limited to, infiltration of adipose and fibrotic tissue into the muscle (65-67), instability in neuromuscular transmission at the neuromuscular junction (12), impaired cross-bridge mechanics and Ca^2+^ handling (13-15, 68, 69), and/or the selective atrophy of the fibers expressing the fast MyHC isoforms (16-18). Our data suggests that the factor(s) responsible must also be able to explain the decrease in the amplitude and the rate of torque development of the involuntary, electrically evoked twitch as these were the two most closely associated variables with the age-related power loss for men and women and in old and very old adults (Fig. 7, Tables 2 & 3). A recent comprehensive mathematical modeling study revealed that simply increasing the proportional area of the muscle composed of MyHC I fibers was able to explain a majority of the altered twitch properties observed with aging (70). In addition, it was recently found that estimates of the thigh lean mass composed of the fast MyHC isoforms were closely associated with the age-related loss in power in men (16). Although we did not measure the fiber type composition of the participants in the present study, the slower rate of twitch torque development, longer relaxation times, and greater impairments in torque-and power-generating capacity at faster velocities observed in the older adults are all consistent with a muscle composed of a lower proportional area of fibers expressing the fast MyHC isoforms. Future studies should examine the association between the loss of power and the selective loss and/or atrophy of the fast MyHC isoforms with aging in men and women across a wide range of ages.

It is important to acknowledge that a portion of the loss in power with aging may also be the result of the decreased physical activity levels and increased sedentary behavior that is commonly observed in older compared with younger adults (71). Indeed, the physical activity of the very old adults in the present study was lower compared with both the young and older adults (Table 1). However, the association between peak power and physical activity was notably weak, with steps per day accounting for only ∼3% of the variance in peak power. This finding is in agreement with a previous study that also found no association between habitual physical activity and peak power output in young and older adults (72). In addition, the older adults in the present study generated 42- 49% lower power than the young, despite having similar physical activity levels, suggesting that the age differences we observed in power were not likely due to differences in physical activity alone. Future studies that include a broader range of older adults from more sedentary to highly active will help answer this important question as lifelong aerobic exercise has been shown to help attenuate at least some of the loss in strength and power with aging (53, 73).

#### Conclusions

Aging was associated with a progressive loss in isometric strength and power output of the knee extensor muscles, with the greatest age differences in power at the faster contraction velocities in both men and women. The rate of decline in absolute maximal isometric strength and power output with age was greater in men than the women but the relative rate of decline did not differ between the sexes. The mechanisms for the loss in lower limb power output of healthy, community dwelling adults also appear to be mostly similar between men and women. Voluntary activation from the nervous system remains near maximal across the lifespan and can account for only a small portion of the age-related loss in power, specifically for older women. We conclude that the accelerated loss in power output relative to muscle mass with aging is determined primarily by factors that alter the contractile function of the muscle in old and very old men and women.

## AUTHOR CONTRIBUTIONS

S.K.H. and C.W.S. conceived and designed the research; A.K., M.A., and C.W.S. performed experiments; D.J.W., A.K., M.A., and C.W.S. analyzed data; D.J.W., S.K.H., and C.W.S. interpreted results of experiments; D.J.W. and C.W.S. prepared figures; D.J.W. and C.W.S. drafted manuscript; D.J.W., S.K.H., and C.W.S. edited and revised manuscript; D.J.W., A.K., M.A., S.K.H. and C.W.S. approved final version of manuscript.

## ACKNOWLEDGMENTS

We thank Bonnie Schlinder-Delap for assistance with scheduling the participants and Drs. Hamidollah Hassanlouei and Minhyuk Kwon for helping with data collection during various phases of the study. We also thank the research participants for their willingness to provide the rigorous efforts necessary to make this study possible.

## Supplementary Material

**Fig. S1.**
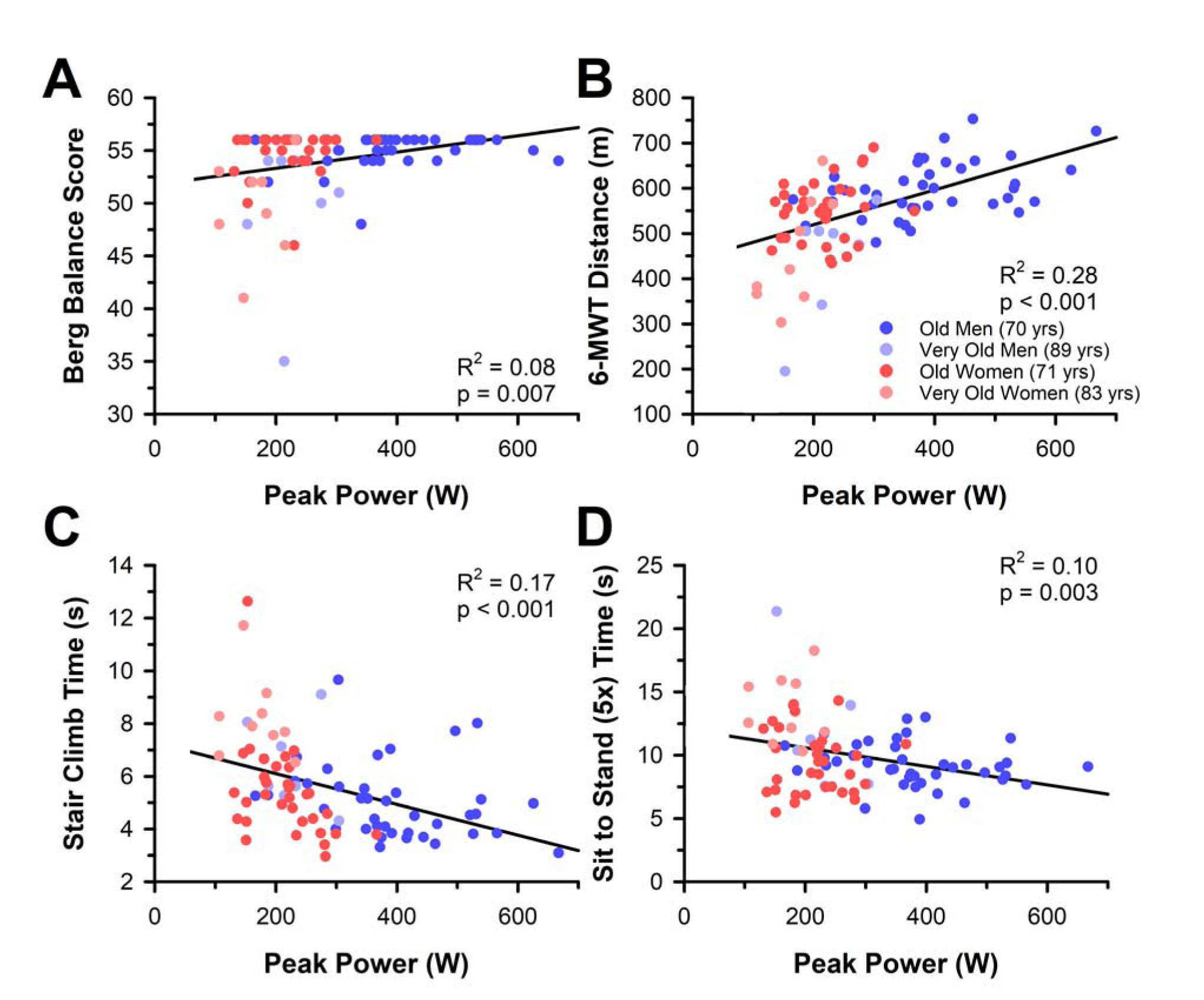
Associations between peak power output and functional performance scores in older men and women. Absolute peak power output was associated with (A) Berg Balance Test scores, (B) the 6-minute walk test distance, (C) stair climb time, and (D) the sit-to-stand time (5x) in the old and very old men and women.

